# Analysis of Gene Expression Heterogeneity Reveals Therapeutic Targets and Novel Regulators of Metastasis

**DOI:** 10.1101/2022.12.16.520816

**Authors:** Dongbo Yang, Christopher Dann, Andrea Valdespino, Lydia Robinson-Mailman, Madeline Henn, Mengje Chen, Gábor Balázsi, Marsha Rich Rosner

## Abstract

Tumor cell heterogeneity has been implicated in metastatic progression of solid tumors such as triple-negative breast cancer (TNBC), leading to resistance and recurrence. We hypothesized that genes with low cell-to-cell transcriptional variability may be effective therapeutic targets, and that analysis of variability may facilitate identification of new metastatic regulators. Here we demonstrate, using single cell RNA sequencing, that the metastasis suppressor Raf Kinase Inhibitory Protein (RKIP) reduced overall transcriptional variability in TNBC xenograft tumors. Focusing on genes with reduced variability in response to RKIP, we identified targetable gene sets such as oxidative phosphorylation and showed that metformin could inhibit RKIP-expressing but not control tumor growth. We also found many regulators of cancer progression including a novel epigenetic metastasis suppressor, KMT5C. These studies demonstrate that a metastatic regulator can alter transcriptional variability in tumors and reveal the importance of genes involved in heterogeneity as potential therapeutic targets and regulators of metastatic progression in cancer.

## Introduction

Metastasis, the dissemination of tumor cells throughout the body, is the primary cause of death from solid tumors such as breast cancer. Characterized by distinct biological states driven by stresses such as oxygen and nutrient loss in the microenvironment, metastatic tumors are largely drug-resistant but the underlying mechanisms are poorly understood. Identifying and characterizing regulators of metastasis and utilizing them to confer sensitivity to treatment is one strategy for developing successful therapeutic intervention. In particular, inhibition of metastasis might promote more homogeneous target expression across cell populations that could enhance therapeutic efficacy.

Multiple lines of evidence suggest that heterogeneity arising from genetic, epigenetic and/or phenotypic sources can be a driver of metastatic progression and drug resistance. For example, heterogeneity in the expression of proteins involved in apoptotic signaling attenuated the apoptotic response of tumor cells to TRAIL (*1*). Similarly, morphological heterogeneity of tumor cells was shown to correlate with increased metastatic potential (*2*). Finally, high intratumoral heterogeneity is strongly correlated with aggressiveness and predicts poor patient outcome (*3*). While heterogeneity is commonly studied at the genetic level by looking at mutations in single cells, emerging evidence suggests that transcriptional heterogeneity is also related to metastatic outcome (*2, 4*).

To identify genes associated with metastasis that could be leveraged as effective drug targets, it is important to assess cell-to-cell variation within the population in addition to mean expression. Previous studies revealed that gene expression variability or noise measured by the coefficient of variation (CV) tends to drop as the inverse of the mean. That is, genes with high mean expression tend to have low noise; and conversely, genes with low mean tend to have high variability (*5–7*). Ηowever, molecular perturbations, epigenetic effects, and feedback or general regulatory pathways can alter the inverse noise-mean interdependence and impact cell function or survival (*5, 6, 8, 9*). For example, drug-based perturbation of genes involved in the DNA repair pathway to amplify noise without significantly changing mean can potentiate transitions in stem cell fate (*5*). Noise-modulating chemicals can also synergize with mean-modulating drugs to promote viral clearance (*10*). In the context of cancer, we have recently shown that increasing the transcriptional noise of a metastasis promoter without altering the mean can promote invasion of tumor cells (*11*). Given that control of both expression mean and variability have direct impact on cell and system-level behavior, we posit that targeting transcriptional variability in addition to mean changes within tumor cell populations could represent an additional strategy to increase treatment efficacy.

In the present study, we sought to test the hypothesis that changes in transcriptional variability can enable identification of metastasis targets and novel regulators. As a possible means of reducing gene expression variability, we ectopically expressed a metastasis suppressor, Raf Kinase Inhibitory Protein (RKIP; encoded by *PEBP1*) in metastatic triple-negative (TNBC) tumor cells (*12*). To measure transcriptional variability, we conducted single cell RNA sequencing (RNA-seq) of metastatic (low RKIP) and non-metastatic (high RKIP) tumor cells and measured both the change in mean and CV of gene expression in cells across the tumor population. We found that RKIP reduces overall transcriptional variability in primary TNBC tumors including genes involved in oxidative phosphorylation. Consistent with more uniform cell-to-cell target gene expression, inhibiting oxidative phosphorylation with the electron transport chain inhibitor metformin suppressed the growth of RKIP-expressing but not control tumors (*13*). Additionally, many regulators of tumor growth and metastasis were less variable in the RKIP-expressing cells including KMT5C, a histone methyltransferase that we show is a novel suppressor of metastasis. Taken together, these results indicate that analysis of gene expression variability along with mean can lead to effective target selection and provides an additional strategy for identifying novel regulators of metastasis.

## Results

### RKIP overexpression reduces transcriptional variability in TNBC

To identify transcriptional differences between metastatic and non-metastatic tumors, we performed single-cell RNA sequencing of human TNBC tumors propagated in a xenograft mouse model. Previously, Rosner and colleagues demonstrated that the TNBC cell line, BM1, lost its metastatic potential upon expression of a metastasis suppressor, RKIP-S153E (*14*). In the present study, following mammary fat pad injection of RKIP-S153E or vector control BM1 cells into athymic nude mice, we dissociated the resulting tumors into single cells and processed the viable human tumor cells for single-cell RNA sequencing (Fig. 1A). To compare transcriptional variability across single cells, it is beneficial to obtain as many sequencing reads as possible from each cell. Therefore, we used a microfluidics-based method that offers more sequencing depth per cell compared to commonly used droplet-based methods (*15*).

**Figure 1.**
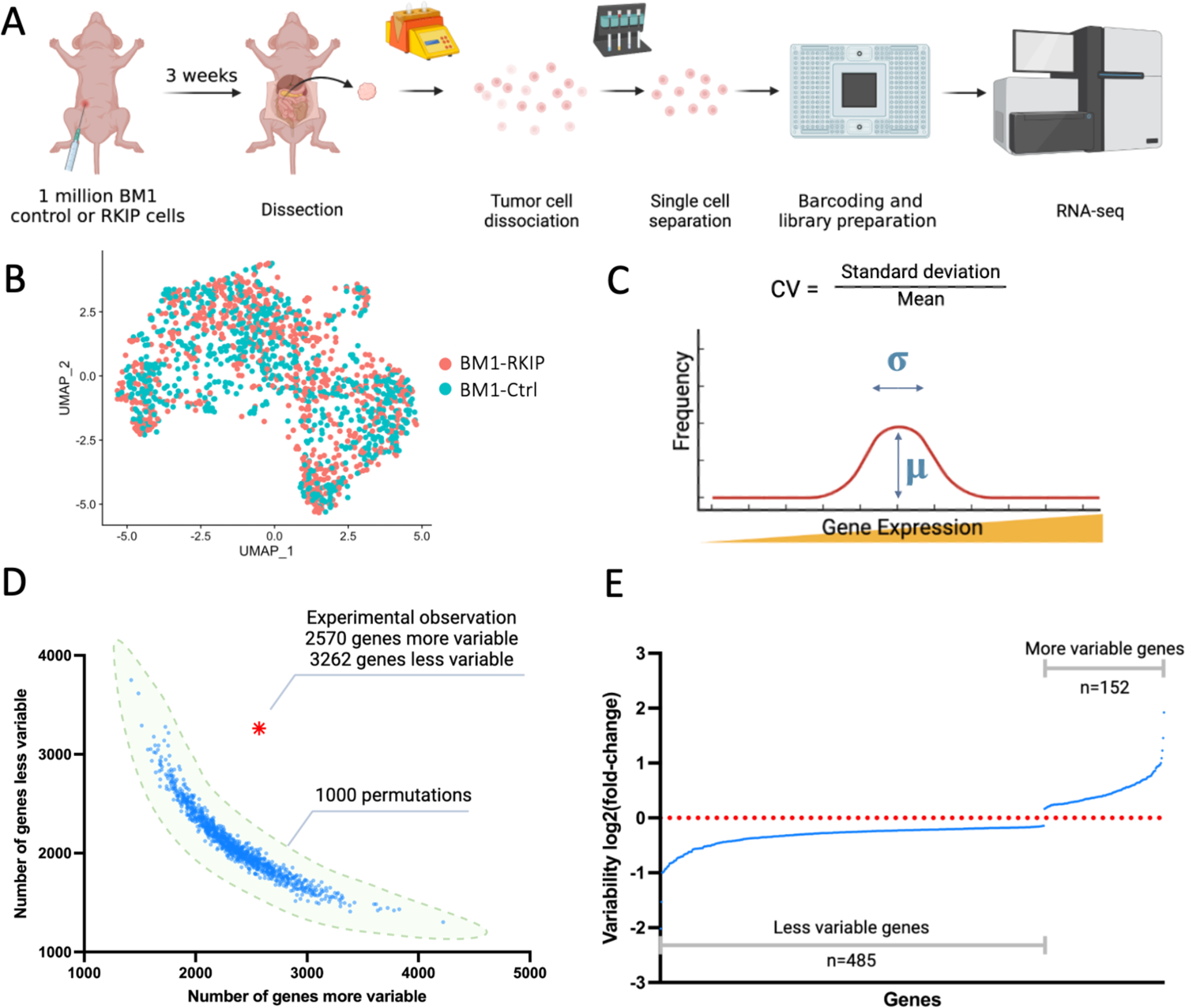
RKIP expression significantly decreases gene expression variability in TNBC xenograft tumors. A) Schematic of the scRNA-seq study. B) UMAP clustering of BM1-Ctrl and BM1-RKIP single cells. C) Calculation of gene expression coefficient of variation from single-cell gene expression mean and standard deviation. D) Number of more variable or less variable genes in BM1-RKIP cells (10% or more difference from control cells) observed by experimental observation compared to 1000 permutations. E) Changes in gene expression variability of genes that were significantly more or less variable in BM1-RKIP cells.

Before analyzing transcriptional variability, we evaluated three independent replicates from each tumor type to control for sequencing quality and depth, reproducibility, cell cycle, and batch effects. Distributions of the number of genes detected per cell were comparable between the metastatic and non-metastatic tumor samples for each set of replicates (Fig. S1A), and the single-cell gene expression profiles were reproducible across the three replicates (Fig. S1B). The distribution of cells within the cell cycle for the two tumor types was also comparable (Fig. S1C). Notably, UMAP clustering could not effectively separate control and RKIP-expressing cells despite their known differences in metastatic potential (Fig. 1B), suggesting that this dimensionality reduction technique is insufficient to distinguish between metastatic and non-metastatic phenotypes.

We first identified genes with differential expression (change in the mean, μ) between control and RKIP-expressing cells in tumors. To account for sequencing noise and gene dropouts due to differences in read counts in single-cell sequencing, we analyzed statistically significant differences in mean gene expression between control and RKIP-expressing tumor cells as previously described (*16*). Of the 15,006 genes characterized, 596 increased in mean and 930 decreased in mean with RKIP overexpression, indicating that transcriptional expression of most genes was unchanged (Table 1). Although the statistical metric based upon the mean is generally used to subset genes and characterize differences between tumor types, we reasoned that a different statistical metric based upon transcriptional variability would generate an alternative gene subset that might identify other potential targets of interest.

**Table 1.**
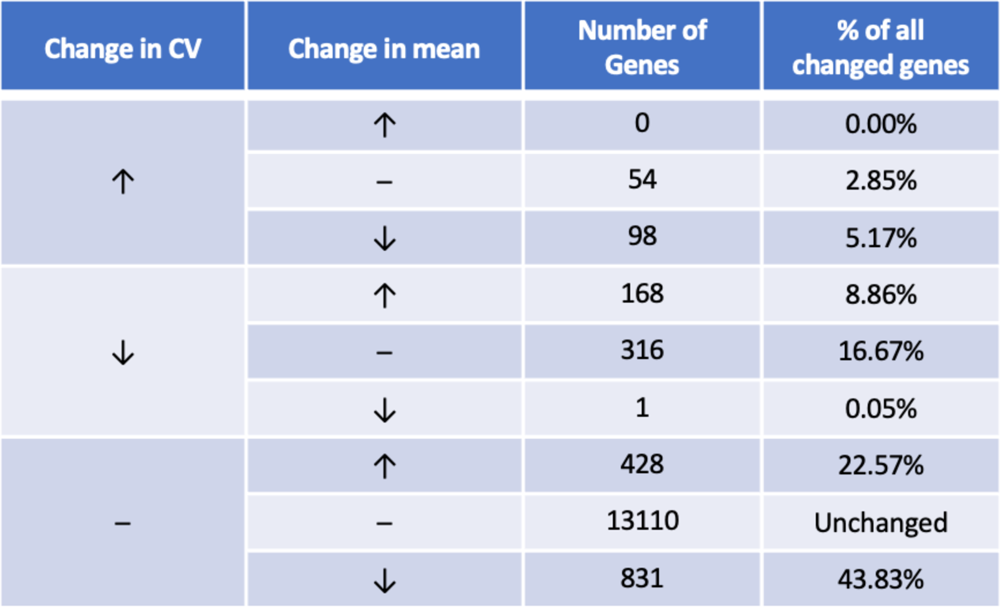
Genes with significant changes in expression mean or CV in BM1-RKIP cells versus BM1-Ctrl cells.

| Change in CV | Change in mean | Number of Genes | % of all changed genes |
| --- | --- | --- | --- |
| ↑ | ↑ | 0 | 0.00% |
|  | – | 54 | 2.85% |
|  | ↓ | 98 | 5.17% |
| ↓ | ↑ | 168 | 8.86% |
|  | – | 316 | 16.67% |
|  | ↓ | 1 | 0.05% |
| – | ↑ | 428 | 22.57% |
|  | – | 13110 | Unchanged |
|  | ↓ | 831 | 43.83% |

To elucidate transcriptional variability among single tumor cells, we calculated the CV (σ/μ where σ refers to the standard deviation; Fig. 1C) for each gene from the RKIP-expressing tumor cells versus control by combining single-cell gene expression results from all three replicates and removing batch effects. Experimental observation showed that RKIP overexpression caused an overall decrease in heterogeneity. To determine whether these observed differences represent true biological differences as opposed to technical artifacts due to the noisiness of single-cell sequencing, we performed 1000 rounds of permutations involving random reassignment of single-cell expression levels to RKIP or control cell types (see Methods) and compared the results of all the permutations to the experimentally observed difference. The observed experimental difference in gene expression variability does not overlap with the distribution of results from 1000 permutations indicating that there are significant differences in gene expression variability between RKIP and control tumor cells (Fig. 1D). Using false discovery rate (FDR)-adjusted P-values derived from the permutation analysis, we found that 485 genes were significantly less variable (more homogeneous) in RKIP cells (including the overexpressed RKIP itself) and 152 genes were more variable (Figs. 1E, S1D).

### RKIP causes changes in gene expression mean and/or variability

Upon analyzing the genes with changes in mean or CV, we identified 20% with CV changes only, 66% with mean changes only, and 14% with both mean and CV changes when comparing control to RKIP-expressing tumor cells (Table 1). There were no genes up-regulated and more variable in RKIP cells, while only one gene was down-regulated and more homogeneous, suggesting that CV and mean rarely increase or decrease together upon RKIP overexpression. Notably, among the genes with increased mean, only a minority (28%) displayed a decrease in CV indicative of more homogeneous expression throughout the population (Table 1; see also Fig. 3A). Among the genes with decreased CV, the majority (65%) had no significant alteration in mean (Table 1).

Constitutive gene transcription is generally considered a Poisson process comprised of random, uncorrelated events of mRNA synthesis that occur at a constant rate (*17*). The mean (μ) and the variance (σ^2^) of a Poisson process are equal. Thus, examining the relationship between the mean and variance of transcriptome data can reveal whether the expression of genes is constitutive or not. That is, a Fano Factor = σ^2^/μ = 1 or a CV^2^= σ^2^/μ^2^=1/μ would be consistent with each gene being expressed with a constant rate, as a Poisson process. On a logarithmic plot, the slope of log_10_(CV^2^) versus log_10_(μ) will then be −1. However, if the rate of transcriptional events fluctuates in time, transcription can depart from a Poisson process in various ways. First, if transcription switches randomly between a high and a low rate, causing transcripts from each gene to arise as intermittent bursts, then the Fano Factor will still be a constant > 1, and the log_10_(CV^2^) versus log_10_(μ) will still have a slope of −1, but will be shifted up. Second, in more complicated scenarios such as feedback regulation (*18, 19*), upstream noise propagation (*20*), or other influences, the Fano Factor may vary and the log_10_(CV^2^) versus log_10_(μ) slope may differ from −1.

To study how gene expression deviates from constitutive, constant-rate synthesis, we plotted log_10_(CV^2^) versus log_10_(μ) for both control and RKIP-expressing cells (suppl Fig. 2). If the slope is −1, then this is consistent with a classic Poisson process or intermittent bursts (characterized by a Fano Factor FF≥1 where FF is σ^2^/μ and σ^2^≥μ) (*8*). For genes from both control and RKIP-expressing cells, the slopes were relatively constant and similar, approaching −1, implying that the transcription of most genes is either constitutive, or it occurs in bursts as it alternates randomly between a high and a low rate. The unchanged slope is not surprising as most genes did not significantly differ in variability (∼96%) upon RKIP overexpression. However, when we plotted only the genes whose mean was unchanged but whose expression was significantly less variable in RKIP-expressing cells, we observed a downward shift for RKIP-expressing cells relative to control cells (Fig. 2A) and a corresponding decrease in FF (Fig. 2B). These results are consistent with the decrease in variability in RKIP cells we observed using the CV-based permutation analysis (Fig. 2B; also see Fig. 1D, E). An exception can occur for genes where the mean increases more than σ, but less than σ^2^, so the CV drops, but the FF increases as we observe for the gene RKIP when overexpressed (Fig. 2C). These results indicate that, for most genes with decreased CV, RKIP expression causes FF reduction for both Poisson and non-Poisson processes.

**Figure 2.**
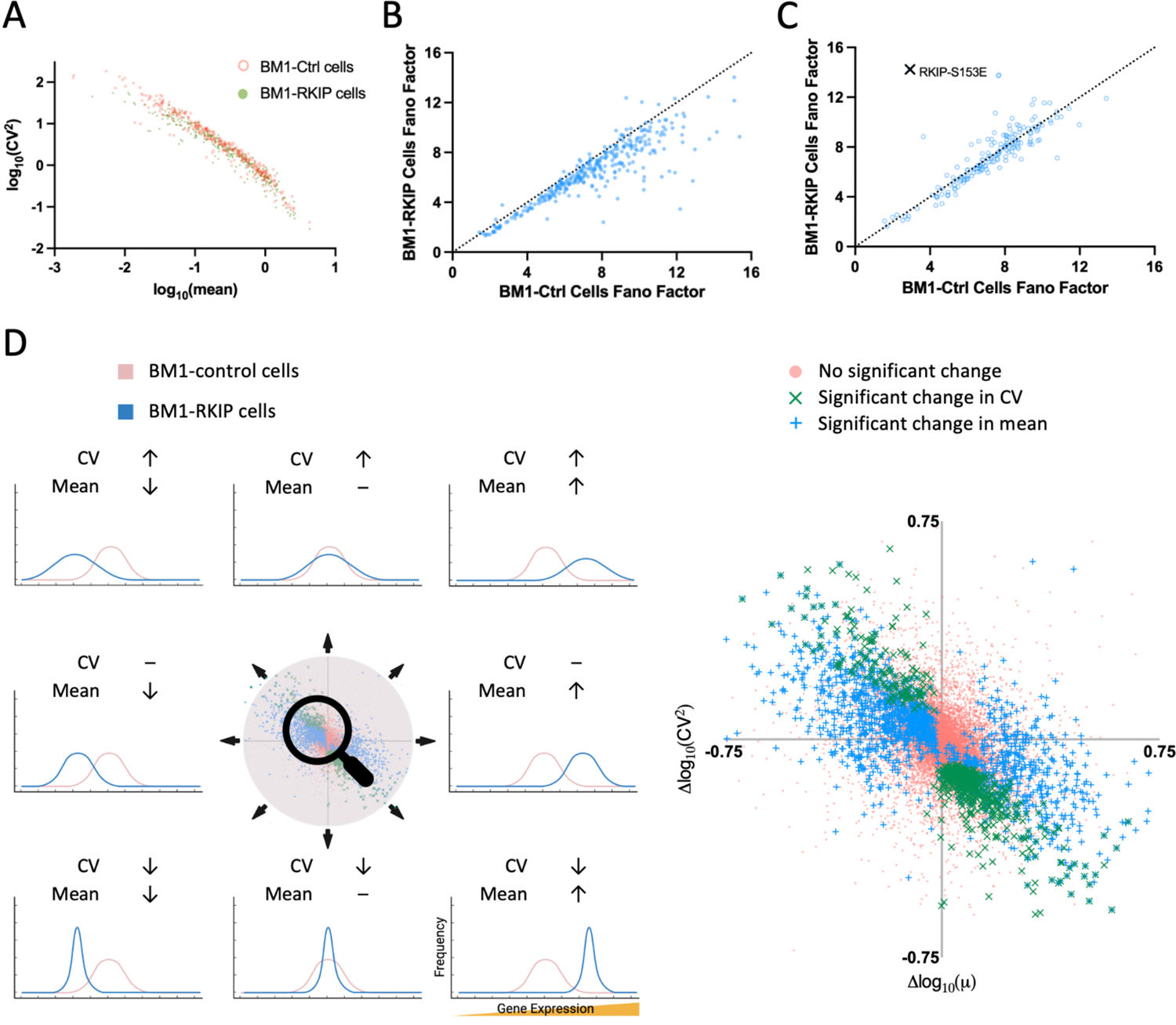
RKIP causes changes in gene expression mean and variability. A) Expression log_10_(CV^2^) versus log_10_(mean) of genes in BM1-Ctrl and BM1-RKIP cells. Genes plotted are those with significant down-regulation in CV without changes in mean in BM1-RKIP cells. B) Fano factor (FF) of genes in BM1-Ctrl cells versus BM1-RKIP cells. Genes plotted are those with significant down-regulation in CV without changes in mean in BM1-RKIP cells. C) FF of genes in BM1-Ctrl cells versus BM1-RKIP cells. Genes plotted are those with significant down-regulation in CV and up-regulation in mean in BM1-RKIP cells. The FF of overexpressed RKIP-S153E in BM1-Ctrl versus BM1-RKIP cells is indicated. D) Left: genes can change in CV and/or mean from BM1-Ctrl to BM1-RKIP cells. Right: enlarged scatter plot of changes in log_10_(CV^2^) and log_10_(mean) for all genes. Genes with significant changes in CV are indicated with “×”, and genes with significant changes in mean are indicated with “+”. Genes with significant changes in both CV and mean are indicated with an overlap of “×” and “+” symbols.

**Figure 3.**
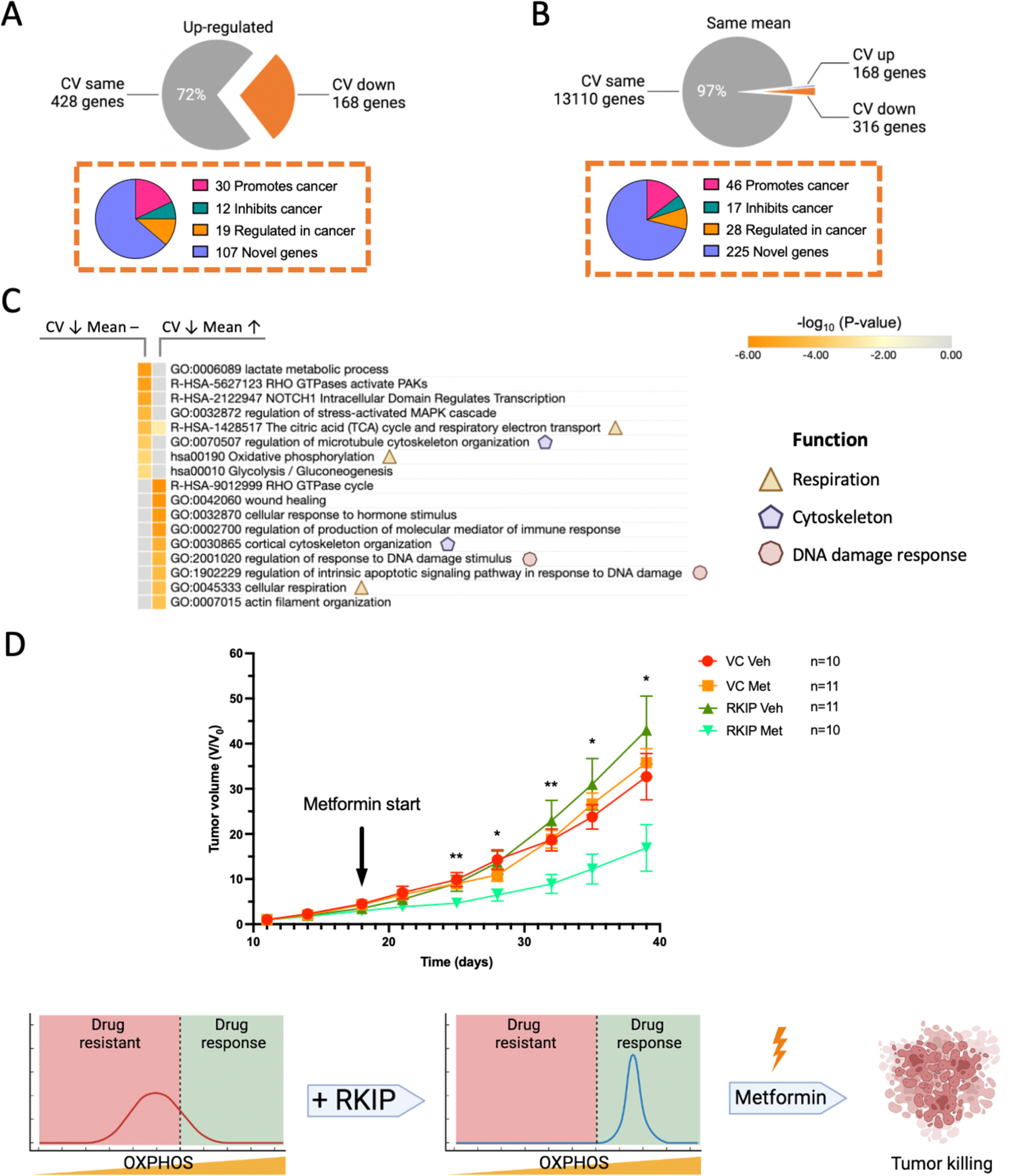
Functions of genes less variable in BM1-RKIP tumor cells. A) Genes with significantly higher mean in BM1-RKIP cells included 168 genes with significantly less CV and 428 genes without CV changes. Genes with less CV contained cancer-relevant genes as shown in the pie chart. B) 3% of all genes without mean changes had significantly higher or lower CV in BM1-RKIP cells. 316 genes less variable in BM1-RKIP cells contained cancer-relevant genes as shown in the pie chart. C) Significantly enriched gene sets from genes less variable in BM1-RKIP cells with either the same mean or up-regulated mean. Common themes of gene sets are highlighted. D) Cells with high RKIP were less variable in OXPHOS and more sensitive to metformin treatment. Mouse tumor volumes are plotted as fold-changes over the initially measured volumes (V/V_0_). Mice were implanted with BM1-Ctrl (VC) or BM1-RKIP (RKIP) tumor cells and were treated with either vehicle (Veh) or metformin (Met). Error bars indicate the standard error of the mean (SEM). Significant differences between VC Veh and RKIP Met were indicated (*: P < 0.05, **: P<0.01; Mann-Whitney test).

We have previously proposed that both gene expression mean and variability changes regulate the transition between metastatic and non-metastatic tumor cells (*21*). Given that CV and/or mean of gene expression can either increase, decrease, or stay the same in cells transitioning to a non-metastatic phenotype, there are nine possible scenarios for the differences ΔCV^2^ and Δmean for each gene (Fig. 2D). Interestingly, Δlog_10_(CV^2^) versus Δlog_10_(mean) followed inverse scaling similar to the log_10_(CV^2^) versus the log_10_(mean) for control and RKIP cells alone. Since we are analyzing differences (Δlog_10_(CV^2^) and Δlog_10_(mean)), this relationship suggests that these changes are constrained in the same way as the original quantities for genes in control and RKIP-expressing cells. This can be explained if the rate of transcription or burst size is increased by a constant factor due to RKIP overexpression (see Supplementary Note).

### Functions of genes with CV changes

Literature searches of genes with CV and mean changes revealed both pro-cancer and anti-cancer genes as well as genes that exhibit context-dependent regulation of cancer (Figs. 3A, B, S3A, Tables S1, S2). Of particular interest are the sets of genes with decreased CV and either increased or unchanged mean that comprise about 25% of all genes with significant changes (Table 1, Fig. S3B). For example, FZD5, the receptor of Wnt5A, contributes to tumor cell proliferation and is suggested to be a drug target in colorectal and pancreatic cancers (Fig. S3B) (*22*). CCND2 is involved in cell cycle regulation (*23*). FGF18 is a fibroblast growth factor that promotes tumor invasion in ovarian cancer (*24*). Genes known to suppress breast cancer are also among the genes that are more homogeneous in RKIP cells. These include FOXP2, a transcription factor that inhibits stemness and metastasis in breast cancer cells (*25*). Suppressor genes are of particular interest with respect to CV as they would not be maximally effective unless they are comparably expressed across the tumor cell population.

We next determined whether the genes with discrete CV and mean changes can be grouped into known cellular functions and processes by performing Metascape analyses for pathway enrichment (*26*). Strikingly, of the 6 configurations of mean/CV changes observed in response to RKIP expression, each corresponded to a discrete set of biological functions (Figs. 3C, S4). Genes more homogeneous in RKIP cells with unchanged or increased means had marked enrichment in oxidative phosphorylation, DNA damage response, and cytoskeleton pathways, while genes more heterogeneous had less defined functions (Figs. 3C, S4). We then checked the seven GO terms highlighted in the combined mean/CV pathway analysis (Fig. 3C) to see if they also appear in CV-only or mean-only analyses (Fig. S5A-B). While the cytoskeleton and DNA damage GO terms show up in all analyses, the combined CV/mean analysis has a significantly stronger P-value and enrichment score (Table 2). Notably, the OXPHOS terms did not even appear as an enriched pathway in the mean-only analysis. Our results suggest that these pathways, while prominently featured in the combined CV/mean analysis, are more likely to be missed using mean analysis alone.

**Table 2.** Enrichment results of GO Terms highlighted in Figure 3C in CV with mean change analysis, CV change-only analysis, and mean change-only analysis. GO terms are color-coded in correspondence to the highlighted functions in Figure 3C (respiration in yellow, cytoskeleton in purple, and DNA damage response in pink). The Analysis column indicates whether the input list of genes belong to CV with mean change analysis, CV change-only analysis, or mean-change-only analysis. The Gene List column indicates the directionality of the genes enriched by Metascape. For each GO term, the Metascape Log_10_(P-value), enrichment score, and Log_10_(Q-value) are shown.

| ID | Pathway | Analysis | Log(P-value) | Enrichment score | Log(Q-value) | Gene List |
| --- | --- | --- | --- | --- | --- | --- |
| hsa00190 | Oxidative phosphorylation | CV & mean | -3.4 | 5.2 | -1.1 | CV down mean same |
|  |  | CV only |  |  |  |  |
|  |  | Mean only |  |  |  |  |
| GO:0045333 | Cellular respiration | CV & mean | -4.1 | 7 | -1.5 | CV down mean up |
|  |  | CV only |  |  |  |  |
|  |  | Mean only |  |  |  |  |
| R-HSA-1428517 | The citric acid (TCA) cycle and respiratory electron transport | CV & mean | -4.1 | 5 | -1.4 | CV down mean same |
|  |  | CV only | -6.1 | 5.1 | -2 | CV down |
|  |  | Mean only |  |  |  |  |
| GO:0030865 | Cortical cytoskeleton organization | CV & mean | -4.8 | 16 | -1.7 | CV down mean up |
|  |  | CV only | -4.4 | 7.6 | -1.3 | CV down |
|  |  | Mean only | -3.2 | 5.7 | -0.49 | Mean up |
| GO:0070507 | Regulation of microtubule cytoskeleton organization | CV & mean | -3.7 | 5.1 | -1.3 | CV down mean same |
|  |  | CV only |  |  |  |  |
|  |  | Mean only |  |  |  |  |
| GO:2001020 | Regulation of response to DNA damage stimulus | CV & mean | -4.4 | 5.5 | -1.6 | CV down mean up |
|  |  | CV only |  |  |  |  |
|  |  | Mean only |  |  |  |  |
| GO:1902229 | Regulation of intrinsic apoptotic signaling pathway in response to DNA damage | CV & mean | -4.3 | 20 | -1.5 | CV down mean up |
|  |  | CV only | -3.5 | 8.5 | -0.95 | CV down |
|  |  | Mean only | -3.3 | 7.5 | -0.51 | Mean up |

We reasoned from these results that the RKIP-expressing cells are more metabolically homogeneous with respect to oxidative phosphorylation and, therefore, more susceptible to respiratory inhibitors like metformin (Fig. 3C). To determine whether RKIP overexpression sensitizes TNBC tumors to metformin treatment, we implanted BM1-CTRL or BM1-RKIP cells in the mouse mammary fat pad, followed by treatment of metformin or vehicle via drinking water. As predicted, mice with BM1-CTRL tumors did not respond to metformin treatment while BM1-RKIP tumors displayed a significant reduction in tumor growth rate and tumor volume (Fig. 3D). These *in vivo* results demonstrate the benefit of analyzing gene expression variability along with expression mean.

Taken together, these results reveal distinct cancer-related genes and functions that become less transcriptionally variable upon losing their cellular metastatic state. Notably, potential druggable targets can be identified by filtering genes with both mean and variability changes.

### CV analyses enable the identification of novel genes that regulate cancer

Most of the genes with altered CV are novel, and their relationships to cancer cannot be ascertained based on current literature. One approach to identifying important regulators of cancer is the analysis of co-expression networks for genes with altered variability. To test this possibility, we performed gene co-expression network analysis for genes with significant CV changes and identified a network of genes with positive correlations in gene expression. The largest network is centered around RPL15. one of the genes with significantly reduced variability but unchanged mean. RPL15 is co-expressed with several cancer-related genes with decreased CV including proliferation markers (MKI67 aka Ki67), metabolic enzymes (LDHA, LDHB), and metastasis suppressors (ARHGDIB) (Figs. 4A, S6). The importance of RPL15 in the context of heterogeneity is supported by a recent study suggesting that more variable RPL15 protein expression is associated with metastasis in breast cancer (*4*).

**Figure 4.**
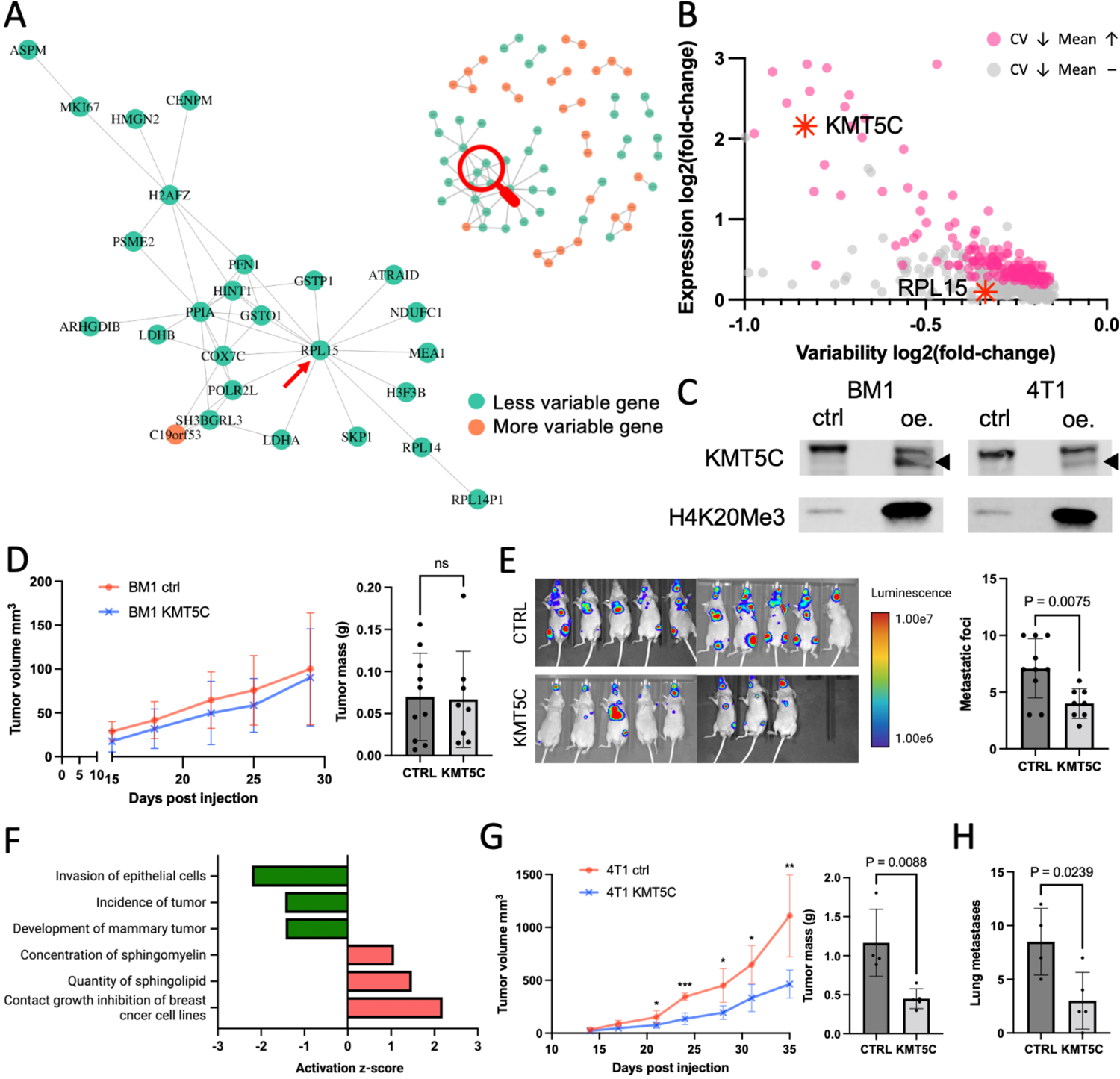
Genes less variable in BM1-RKIP cells include regulators of metastasis. A) Gene co-expression network within the BM1-RKIP cells with the largest subnetwork enlarged. B) KMT5C and RPL15 were less variable in BM1-RKIP cells. C) BM1 and 4T1 cells with stable KMT5C expression also had increased trimethylation of H4K20. D) Mouse BM1 control or KMT5C tumor volumes and tumor mass at dissection. E) Left: luminescence measurement of mice 3 weeks post intracardiac injections of BM1 ctrl (CTRL) or BM1 KMT5C (KMT5C) cells. Right: number of metastatic foci counted in each mouse. F) Activity changes of disease and biological functions in BM1 KMT5C tumors versus control tumors, predicted by Ingenuity Pathway Analysis (IPA). G) Mouse tumor volumes of 4T1 control or KMT5C and tumor mass at dissection. H) Number of metastases on the surface of lungs from BALB/c mice with 4T1 ctrl or KMT5C primary tumors.

An alternative approach for identifying novel regulators of metastasis is to characterize genes with decreased CV and increased mean. KMT5C is among the top six genes with most decreased variability and was significantly up-regulated in RKIP-expressing tumor cells (Figs. S3A, 4B). KMT5C tri-methylates H4K20Me3 and acts as an epigenetic repressor (*27*). However, its role in breast cancer progression has not been addressed, and there are conflicting reports for other cancer types (*28–30*). To test whether KMT5C regulates metastasis, we generated TNBC cells with stable over-expression of KMT5C and confirmed the increased tri-methylation at histone H4K20 (Figs. 4C, S7A). Cells with or without KMT5C overexpression showed comparable proliferation *in vitro* (Fig. S7B). Similarly, xenograft tumors in athymic nude mice with or without KMT5C overexpression did not exhibit differences in primary tumor growth (Fig. 4D) but did show a marked increase in H4K20Me3 staining (Fig. S8A). By contrast, intracardiac injections of KMT5C-expressing BM1 cells resulted in significantly fewer metastatic colonies compared to the control cells, consistent with a role for KMT5C as a metastasis suppressor (Fig. 4E). RNA sequencing of the primary xenograft tumors identified 814 genes down-regulated and 769 genes up-regulated in KMT5C-expressing tumors (Fig. S8B). Disease and biological functions analyses of the differentially expressed genes identified changes in metastasis-related functions in KMT5C-overexpressing tumors (Fig. 4F). These results were validated using a syngeneic mouse model showing that KMT5C suppressed primary tumor growth, possibly due to the intact immune system, as well as spontaneous metastases to the lung (Fig. 4G-H). Together, these results suggest that KMT5C functions as a metastasis and context-dependent tumor suppressor for TNBC.

### KMT5C is a clinically relevant regulator of metastasis in breast cancers

We next analyzed expression levels of KMT5C in cancer patients. We previously identified a group of invasion genes (INV) suppressed by RKIP or driven by BACH1, an RKIP-inhibited transcription factor that promotes metastasis in breast cancer (*31*). We also identified genes in the electron transport chain (OXPHOS) that are up-regulated by RKIP expression (see Fig. 3C) or BACH1 depletion in TNBC cells (*13*). Analysis of the TCGA database across multiple solid tumor types for expression of these genes shows a significant negative correlation between KMT5C and INV gene expression and a positive correlation of KMT5C to OXPHOS gene expression (Figs. 5A, S9). Similarly, within individual patient samples, both breast and pancreatic tumors with higher KMT5C expression generally express lower levels of INV genes and higher levels of OXPHOS genes (Figs. 5B, S10A, B). We then compared KMT5C to BACH1 expression in patients and observed a negative correlation. In addition, we stratified breast cancer patients by their relative expression levels of either KMT5C, RKIP, or BACH1 and performed differential expression analyses within each patient group (Fig. S11 A-C). Both RKIP and KMT5C are associated with an overlapping set of canonical pathways in a manner that suppresses cancer progression (Fig. 5C). By contrast, the same pathways are associated in the opposite manner with BACH1.

**Figure 5.**
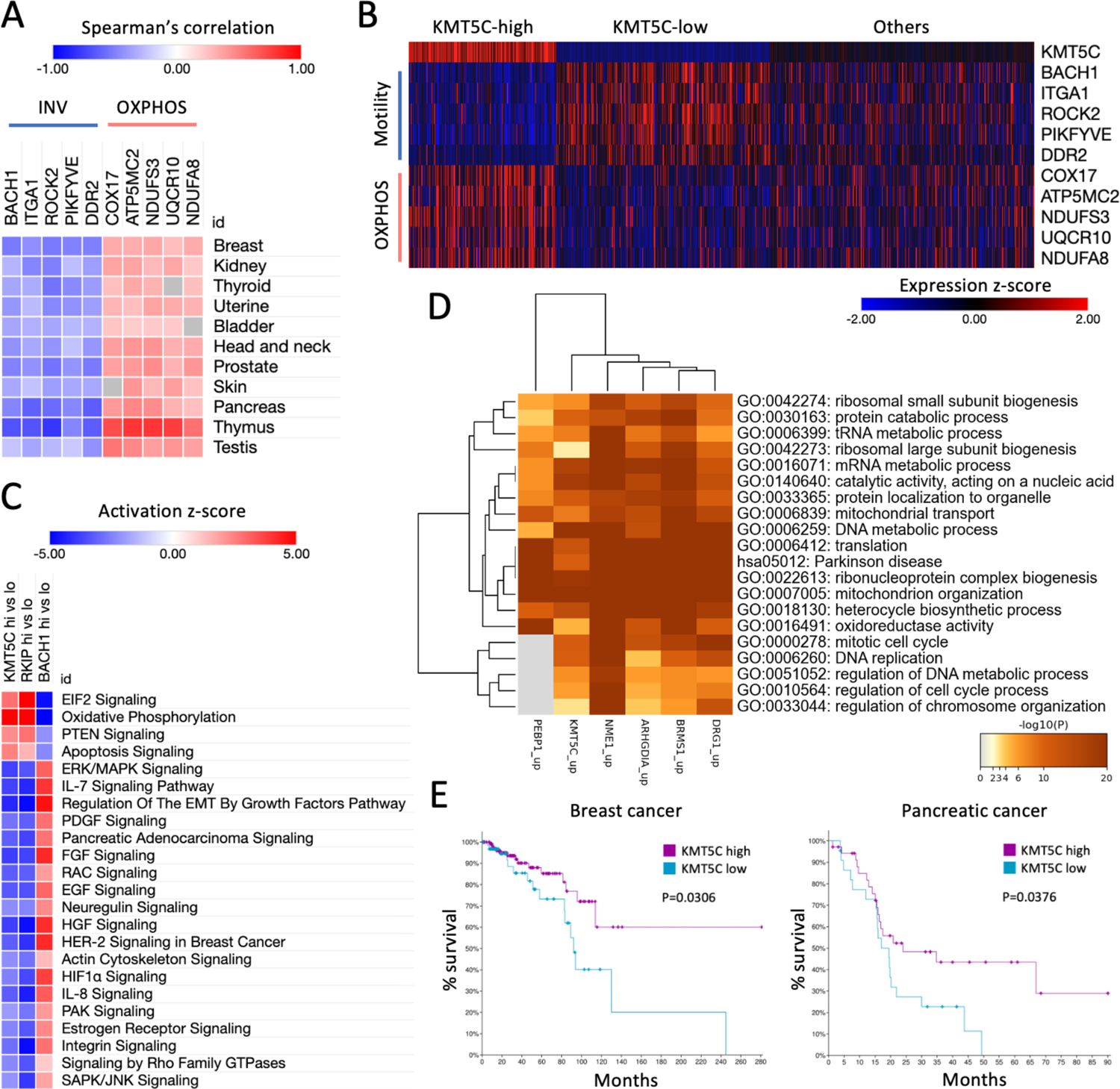
KMT5C expression correlates with better survival and less metastasis in patients. A) Spearman’s correlation between KMT5C and genes related to metastasis in cancer patients. B) Relative expression levels of metastasis-related genes (INV/OXPHOS) in breast cancer patients with higher or lower levels of KMT5C. C) Predicted activation of canonical pathways in TNBC patients with high versus low levels of KMT5C, RKIP, or BACH1. D) Significantly enriched gene sets from genes positively correlated with PEBP1, KMT5C, NME1, ARHGDIA, BRMS1, or DRG1 in breast cancer patients. E) Kaplan-Meier plots of breast cancer (left) or pancreatic cancer (right) patients with high or low levels of KMT5C.

**Figure 6.**
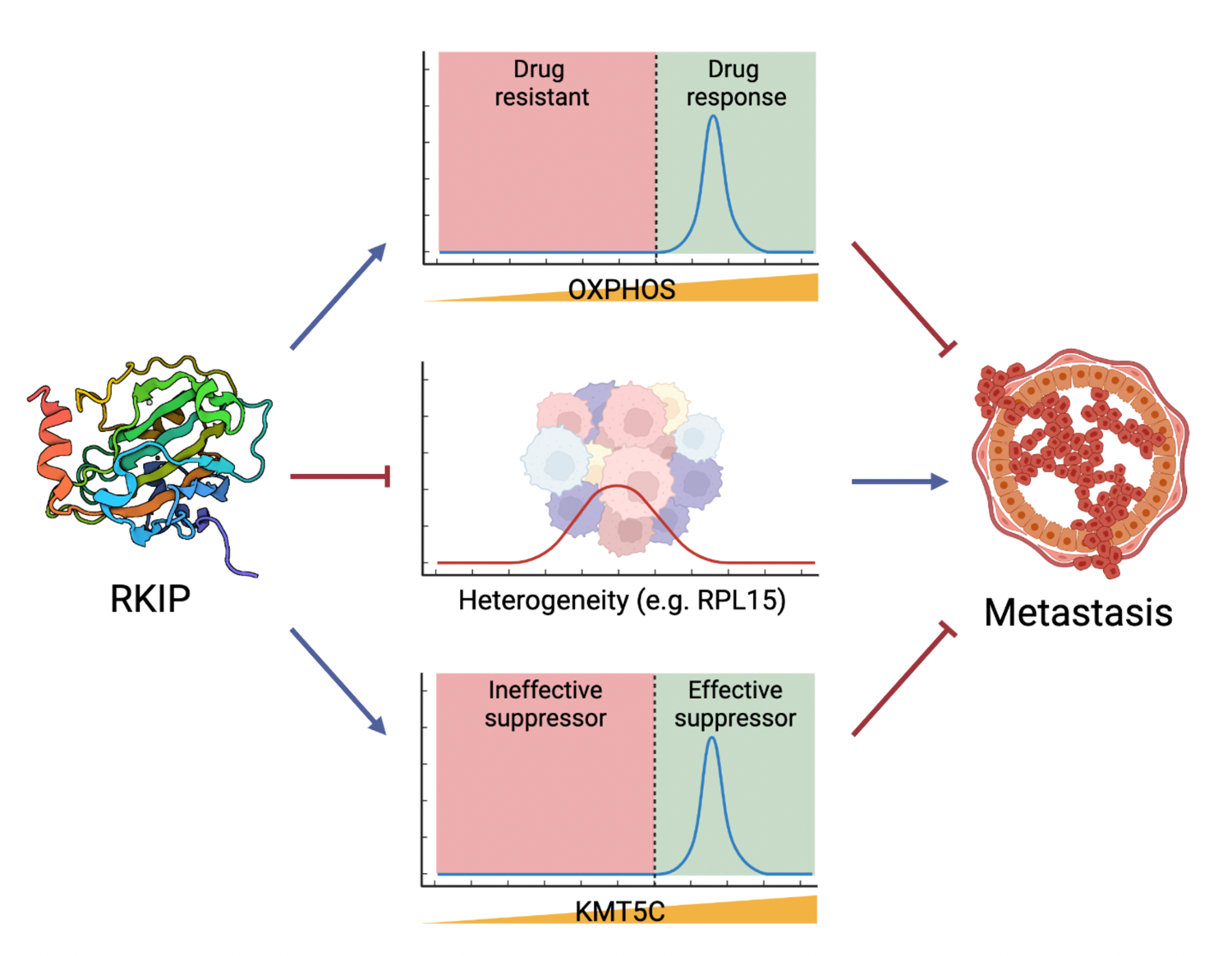
RKIP inhibits overall gene expression variability in non-metastatic tumor cells and reduces variability while up-regulating KMT5C and OXPHOS genes.

We also compared KMT5C to other previously identified metastasis suppressors using the TCGA database (*32, 33*). In breast cancer patients, KMT5C and four known suppressors of metastases (NME1, BRMS1, ARHGDIA, DRG1) are positively or negatively correlated to a similar set of gene functions (Figs. 5D, S12), suggesting that they act in a similar manner. When analyzed alone, KMT5C expression is positively correlated to processes such as translation and aerobic respiration and negatively correlated to several cancer-promoting signaling pathways in addition to invasion and motility (Fig. S13). Finally, in breast and pancreatic cancer patients, higher KMT5C expression is correlated with significantly better overall survival (Fig. 5E). Together, these patient tumor analyses further support the experimental data suggesting that KMT5C, like its upstream regulator RKIP, also functions as a metastasis suppressor and is associated with improved survival in breast and possibly other cancers.

## Discussion

In this study, we use the CV as an additional metric along with mean for comparing differences in transcriptional variability between metastatic and non-metastatic tumor states and to identify effective therapeutic targets as well as novel regulators of cancer progression and metastasis. Our results showed that the metastasis suppressor RKIP can reduce the transcriptional variability of TNBC tumors. Analysis of functional pathways with decreased CV and unchanged or increased mean reveal potential drug targets such as oxidative phosphorylation that enhance tumor sensitivity to treatment. We also report that KMT5C, one of the novel genes up-regulated and more homogeneous in RKIP-overexpressing cells, is a histone modifier with metastatic suppressor function in TNBC.

Previous studies on tumor heterogeneity focused primarily on genetic heterogeneity extrapolated from single-cell mutations and copy number variations or subclonal populations inferred from bulk sequencing data (*34*). By contrast, the CV or similar measures such as the Gini index are only beginning to gain traction as an approach to studying tumor cell heterogeneity at the transcriptional level (*35*). For example, CV was used to quantify cancer cells’ variable apoptotic response to killing (*1*) or metastatic potential (*2*). More recently, the CV method was employed to study transcriptional noise during DNA damage repair, demonstrating the usefulness of this method outside of cancer research (*5, 7*). CV can be used on many types of datasets other than single-cell RNA sequencing or cell-to-cell phenotypic variability. For example, CVs can also be determined from flow cytometry and spatial datasets. With the rise of spatial techniques, we predict this approach can also benefit emerging techniques, including cyclic immunofluorescence and spatial transcriptomics.

As CV can reflect many sources of noise, we also analyzed the Fano factor and demonstrated that RKIP reduces FF for most genes with decreased CV. An alternative metric to measure variability, the Fano factor is commonly used to study deviations from a Poisson process. Unlike CV, which is not affected by scaling during gene expression normalization (both the standard deviation and the mean are scaled by the same scaling factor, thus canceling out), the Fano factor has the same units as the mean. Hence, its value is influenced by the normalization of gene expression, making it difficult to compare Fano factors of genes between different studies (*5*). However, the relative change in the Fano factor between experimental conditions that we observed indicates deviations from a Poisson process, with possible causes including altered bursting, feedback regulation, or noise modulators (*5, 6, 8*).

The mechanism by which RKIP perturbs the Fano factors of many genes is not yet known. Previous studies have shown that changes in DNA damage repair genes can lead to increased noise without changing mean gene expression (*5*). Thus, one mechanism by which RKIP might decrease variability is via DNA repair. Similarly, changes in epigenetic processes could alter transcription. One gene by which this might occur is the RKIP-regulated KMT5C. Future studies will be required to elucidate the precise mechanisms by which RKIP regulates transcriptional variability.

The RKIP-overexpressing tumor cells had more homogeneous expression for genes participating in oxidative phosphorylation, DNA damage response, and cytoskeleton. These findings prompted us to predict that tumors with high RKIP could be sensitive to treatment by inhibitors against these pathways. We tested one of these pathways, OXPHOS, using the well-studied electron transport chain inhibitor metformin, and verified this prediction. Of note, this result is consistent with our previous report that loss of BACH1, a pro-metastatic transcription factor that is repressed by RKIP (*31, 36*), switches the metabolic state of TNBC tumors from aerobic glycolysis to largely oxidative phosphorylation, resulting in therapeutic sensitivity to metformin treatment (*13*). The present finding provides an additional explanation of our previous metformin study suggesting that single cells depleted of BACH1 have more homogeneous expression of oxidative phosphorylation genes across the tumor population. These therapeutically important pathways may not be easily identified using expression mean analysis alone as in the present study, thus demonstrating the effectiveness of our combinatory approach.

We predict that targeting genes with lower CV is more effective than targeting genes without changes in CV. However, testing this hypothesis in tumors is difficult as, currently, it is technically challenging to reduce CV without also changing mean expression outside of tightly controlled tissue culture conditions (*37*). It should also be noted that a small number of genes became more heterogeneous in RKIP tumors. The significance of genes with higher CVs in non-metastatic tumors remains to be determined. Nevertheless, our study reveals that a suppressor of metastasis overall reduces transcriptional variability in tumor cells, and analyzing genes with decreased CV in combination with mean changes can lead to the identification of clinically important targets and novel regulators of metastasis.

## Materials and methods

### Cell lines

The human TNBC cell line BM1 (also known as MDA-MB-231-1833, BoM1), the mouse TNBC cell line 4T1, and the human embryonic kidney epithelial cell line 293T were used in our studies. 4T1 and 293T cells were purchased from American Type Culture Collection (ATCC). BM1 cells were generated by Massagué and colleagues (*38*). BM1-vector control (BM1-VC) and BM1-RKIP-S153E (BM1-RKIP) cells were generated from wild-type BM1 by Dangi-Garimella and colleagues (*14*). Luciferase-expressing BM1 cells (BM1-luc) cell lines were generated from wild-type BM1 by Yesilkanal and colleagues (*31*). Stable BM1 and 4T1 cells with vector control or KMT5C overexpression were generated through lentiviral transduction following established protocols (*31*). The cloning method for the KMT5C overexpression lentiviral vector is described below. BM1 and 293T cells were cultured in DMEM media (Gibco, 11965-118) with 10% fetal bovine serum (FBS, Avantor, 97068-085) and 1% penicillin-streptomycin (Gibco, 15140-122). 4T1 cells were cultured in RPMI media (Cytiva, SH30027.01) with 10% FBS and 1% penicillin-streptomycin. Cells were grown in 5% CO_2_ incubators at 37 °C. All cell lines were authenticated by short tandem repeat (STR) analysis (IDEXX BioAnalytics) and mycoplasma detection (Lonza, LT07-218) routinely.

### Generation of KMT5C overexpression lentiviral vector

The human KMT5C (NM_032701.4) open reading frame (ORF) was cloned from the pCMV6-Entry expression vector (Origene, RC203881) into the pLV-EF1a-IRES-Hygro lentiviral backbone (Addgene, 85134) with restriction enzymes BamHI-HF (New England BioLabs, R3136S) and MluI-HF (New England BioLabs, R3198S). A stop codon was then added to the end of the ORF through site-directed mutagenesis (New England BioLabs, E0554S) using the following forward and reverse primers: TAACGCGTGAATTCCTCGAG; CAGCTCTTCACCGCCGAC. The resulting pLV-EF1a-IRES-Hygro-KMT5C plasmid was confirmed by Sanger sequencing at the University of Chicago Comprehensive Cancer Center DNA Sequencing and Genotyping Facility. These primers were used: TCAAGCCTCAGACAGTGGTTC; TTGTGCCTGCAGATGGGAACGC.

### Mouse studies

Mice were acquired and housed by the Animal Resources Center at the University of Chicago. All mouse studies were conducted following protocols approved by the Institutional Animal Care and Use Committee. For primary tumor growth experiments, tumor growth was monitored over time by caliper measurements of the width and lengths of tumors using the following formula for tumor volume calculations (*13*):

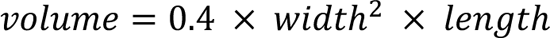

The mice were sacrificed at the experiment endpoint or when the tumor volume reached 1 cm^3^, whichever came first.

For single-cell RNA-seq studies, 1 × 10^6^ BM1-VC or BM1-RKIP cells were injected orthotopically into the mammary fat pad of athymic nude mice (Harlan Envigo). The mice were sacrificed three weeks after tumor cell implantation. Primary tumors were collected for single-cell isolation.

For the metformin treatment study, 1 × 10^6^ BM1-VC or BM1-RKIP cells were injected orthotopically into the mammary fat pad of athymic nude mice (Charles River). When tumors reached 20-30 mm^3^ in volume, mice were randomized into groups for treatment with metformin or vehicle. Metformin (MilliporeSigma, PHR1084) was provided in drinking water *ad libitum* at 300 mg kg^-1^ day^-1^. Tumor growth was monitored until the experiment endpoint was reached.

For BM1 KMT5C primary tumor growth experiments, 1 × 10^6^ BM1-VC or BM1-KMT5C cells were injected orthotopically into the mammary fat pad of athymic nude mice (Charles River). The mice were euthanized four weeks post tumor cell implantation. Primary tumors were collected for RNA sequencing and immunohistochemistry (IHC) analyses.

For BM1 KMT5C colonization assays, 1 × 10^5^ BM1-luc cells with or without KMT5C overexpression were injected into the left ventricle of the heart of athymic nude mice (Charles River) for systemic distribution of tumor cells. Three weeks post-injection, mice were injected with freshly-prepared luciferin solution (Goldbio, LUCK-1G) with a concentration of 15 mg/ml in PBS, and each mouse received 150 mg luciferin/kg of body weight. Five minutes after luciferin injection, tumor burden was measured as bioluminescence via the IVS Spectrum Imaging System (PerkinElmer), and up to five mice were measured at a time. For each group of mice, bioluminescence (radiance, p/sec/cm^2^/sr) was measured continuously until the maximum signal intensity was reached. The mice were euthanized after imaging. The number of metastatic foci for each mouse was determined by counting the number of centroids from the peak radiance measurement. The same setting and radiance scale (1.00e6 – 1.00e7 p/sec/cm^2^/sr) were used to evaluate all the mice.

For 4T1 KMT5C primary tumor growth and spontaneous metastasis assays, 1 × 10^4^ 4T1-VC or 4T1-KMT5C cells were injected orthotopically into the mammary fat pad of BALB/c mice (Charles River). The mice were euthanized four weeks after tumor cell implantation. Primary tumors were collected for immunohistochemistry (IHC) analyses. Lungs were perfused with PBS injected horizontally through the trachea, removed from the thoracic cavity, and then fixed with Bouin’s Solution on (Ricca Chemical, RC1120-16). Spontaneous metastases were measured by counting the number of surface metastases on the lungs following fixation.

### Preparation of single cells for RNA sequencing

BM1 cells from the primary tumors were isolated through tumor dissociation, mouse cell depletion, and dead cell removal following the manufacturer’s protocols (Miltenyi Biotec, 130-095-929, 130-104-694, 130-090-101). Suspensions of single cells were processed on a Fluidigm C1 instrument using a C1 Single-Cell mRNA Seq HT IFC and Reagent Kit v2 (Fluidigm, 101-4981, 101-3743). cDNA synthesis, barcoding, library preparation, and quality check were performed following the manufacturer’s protocol (Fluidigm, PN 101-4964 I1). All single-cell RNA libraries were sequenced by an Illumina HiSeq 4000 to generate paired-end 100bp reads at the University of Chicago Genomics Facility following standard protocols. Each library consisted of single cells from one BM1-VC tumor sample and one BM1-RKIP tumor sample.

### Differential expression analyses of single-cell RNA-seq

The RNA sequencing data were demultiplexed using the manufacturer-supplied API script to generate individual FASTQ files for single cells based on the barcodes used (Fluidigm, mRNASeqHT_demultiplex.pl). The reads were mapped to the human genome (UCSC hg38 with GENCODE annotations) using RNA STAR (*39, 40*). The resulting mapped reads from every single cell were counted by featureCounts to generate per-gene read counts (*41*). Gene expression data from three biological replicates were combined, and the batch effect was corrected via the “BatchCorrectedCounts” function in the CountClust package by specifying the replicate number as the variable to regress out (*42*). Gene expression data were analyzed using Seurat for quality control, count normalization, UMAP clustering, and cell cycle analyses (*43*). Specifically, barcodes with exceptionally low or high total read counts were filtered with the following parameters for each replicate: for replicate 1, nCount_RNA between 2 × 10^4^ and 6 × 10^5^; replicate 2, nCount_RNA between 2 × 10^4^ and 3 × 10^5^; replicate 3, nCount_RNA between 2 × 10^4^ and 5 × 10^5^. Mitochondrial reads were removed prior to downstream analyses. Genes expressed in fewer than 10 RKIP cells or VC cells were filtered. Gene expression data were normalized using the NormalizeData function with the LogNormalize method and a scale factor of 10000. Differential expression analysis was performed using the SCDE package to counter dropout and amplification biases (*44*). The gene expression log_2_(fold-change) of RKIP tumor cells over VC tumor cells was determined from the “scde.expression.difference” function using the “maximum likelihood estimate” result. Genes with significant differential expression were determined with a cutoff of +/- 0.5 on false discovery rate (FDR)-corrected Z-scores.

### Coefficient of variation (CV) analyses

The formula for calculating gene expression CV:

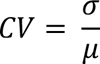

Where σ is the standard deviation of normalized gene expression of a gene across a set of single cells, and μ is the mean of normalized gene expression of the same gene across the same set of single cells. For every gene in the normalized expression dataset, a CV was calculated for each cell type (control and RKIP). The difference in the CV of each gene between control and RKIP tumor cells is determined by calculating the fold-change:

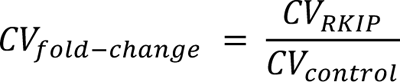

Genes with significant differences in CV were determined through permutations testing, during which 1000 rounds of permutations were performed on the gene expression data. Then, the experimentally observed CV fold-change for each gene was compared against the 1000 permutated CV results to determine the likelihood of obtaining the experimental result by chance, defined by the number of times the permutated result met or exceeded the experimental CV result over 1000 permutations. Genes significantly more or less variable in RKIP cells over control cells were determined with an FDR-adjusted P-value cutoff of 0.05.

### Fano factor of genes in control versus RKIP cells

The formula for calculating gene expression Fano factor (FF):

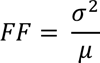

Where σ is the standard deviation of normalized gene expression of a gene across a set of single cells, and μ is the mean of normalized gene expression of the same gene across the same set of single cells. For every gene in the normalized expression dataset, an FF was calculated for each cell type (control and RKIP). However, due to the scaling introduced during count normalization by Seurat, FF will scale accordingly when the standard formula is used to calculate FF (*5*). Therefore, FF is then re-scaled to neutralize the effect of such scaling prior to downstream analyses:

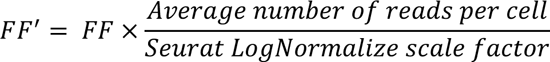

Where the average number of reads per cell is 103543 for RKIP cells and 97545 for control cells, and 10000 was the scale factor applied during the Seurat normalization.

### Network analysis

The gene co-expression network of BM1-RKIP cells was generated from expression correlations between significantly variable genes. Specifically, a Spearman correlations matrix was calculated to determine expression correlations among the genes with significant differences in CV. Normalized expression levels from all BM1-RKIP cells were used to calculate the correlations matrix. The R package “igraph” was used to generate a gene co-expression network from the correlations matrix using the “graph_from_adjacency_matrix” function (weighted and undirected) (*45*). Genes with significantly higher or lower CV were designated as the network nodes, and the edges represented positively correlated gene pairs with correlation coefficients of at least 0.3. Nodes without peripheral connections were excluded from the network. The largest interconnected component of the whole network was extracted using the “decompose.graph” function and plotted again.

### Gene functional enrichment analyses

Metascape was used to determine functions and pathways from lists of genes (*26*). Methods for using Metascape were previously described (*46, 47*). Metascape analyses in the paper were based on the following lists of genes as inputs: genes with or without significant differences in expression CV and/or mean between BM1-RKIP and BM1-VC cells, according to the mouse tumor single-cell RNA-seq studies; genes that correlated with expression levels of KMT5C, NME1, PEBP1, BRMS1, ARHGDIA, or DRG1 in breast cancer patients, accessed through cBioPortal (TCGA, Firehose Legacy, Breast Invasive Carcinoma, mRNA expression RNA Seq V2 RSEM) (*33, 48*). The following cutoffs were used to determine significance: genes with significant CV differences were determined with an FDR-adjusted P-value cutoff of 0.05 from the permutations analyses; genes with significant differential expression were determined with an FDR-adjusted Z-score cutoff of +/- 0.5 from the single-cell differential expression analyses; genes that positively or negatively correlate with the genes of interest in breast cancer patients were selected with a Spearman’s correlation coefficient cutoff of +/- 0.3. The significance of gene set enrichment was determined by P-value < 0.01. Metascape queried the following databases for analyses: GO Molecular Functions, GO Biological Processes, Reactome Gene Sets, and KEGG Pathway (*49–51*).

### Cell proliferation assays

Two thousand tumor cells were seeded per well in a 96-well tissue culture plate. For each cell type, ten wells were seeded. The tissue culture plate was incubated for 24 hours before transferring into an IncuCyte S3 (Sartorius) with the same incubation conditions. Cells were monitored over time by scanning the plate with a 10X phase-contrast objective for up to 7 days. Four images were acquired per well every 4 hours and analyzed by the IncuCyte 2020B software (Sartorius) to determine cell confluence. The mean and standard deviation of percent cell confluence across ten wells were calculated for each cell type at each time point. Proliferation curves were plotted by the IncuCyte software, and error bars represent standard deviations.

### Protein isolation and western blots

Cultured cells were washed with cold PBS and lysed on ice in RIPA buffer with protease inhibitors (Santa Cruz, sc-24948; EMD Millipore, 539134). Samples were then sonicated three times for 10 seconds each at 25-30% power and centrifuged at max speed for 15 min at 4°C. The supernatant was collected, and the protein concentration was quantified using the Bradford assay (Bio-rad, 5000202). Samples were then boiled in 4X Laemmli sample buffer (Bio-rad, 1610747) at 98°C for 5 mins before storage. SDS-PAGE was performed using 20 – 25 μg protein from each sample with a prestained protein ladder (ThermoFisher, 26616). Western blotting was performed using antibodies against KMT5C (AbClonal, A16235, ∼52 kDa) or H4K20Me3 (Cell Signaling, 5737S, ∼11 kDa). Blots were stained for total protein using the REVERT 700 Total Protein Stain Kit (LI-COR, 926-11016) for control. Following primary antibody incubation, blots were then incubated with fluorescent or HRP-conjugated secondary antibodies depending on the target (LI-COR, 926-3221 for KMT5C blots; MilliporeSigma, AP187P for H4K20Me3 blots). Before imaging, blots with HRP secondary antibodies were treated with chemiluminescent HRP substrate (MilliporeSigma, WBKLS0500). Images were acquired on the LI-COR Fc Imaging System and quantified using Empiria Studio Software (LI-COR) with total protein normalization.

### Immunohistochemistry

Mouse tumors were fixed in 10% formalin solution (Azer Scientific, NBF-4-G), then stored in 70% ethanol. Paraffin embedding, cross-sectioning, and IHC staining were performed by the University of Chicago Human Tissue Resource Center following standard protocols. One slide of each tumor sample was stained by the hematoxylin and eosin stain. For H4K20Me3, slides were stained using the Recombinant Anti-Histone H4 (tri methyl K20) antibody (Abcam, ab177190) at 1:4000 dilution. Stained slides were scanned by a CRi Pannoramic SCAN 40x Whole Slide Scanner at the Integrated Light Microscopy Core, then analyzed with CaseViewer software (3DHISTECH).

### Bulk tumor RNA-seq

Five BM1-Ctrl and five BM1-KMT5C tumors were collected from ten athymic nude mice. Fresh tumors were placed inside M Tubes (Miltenyi Biotec, 130-093-236), then homogenized with TRI Reagent (Zymo Research, R2050-1) by the gentleMACS Dissociator (Miltenyi Biotec 130-093-235). Total RNA extraction with genomic DNA removal was performed using the Direct-zol RNA MiniPrep Plus kit (Zymo Research, R2072). The University of Chicago Genomics Facility performed RNA sample quality checks, library construction, and sequencing following standard protocols. All ten samples were sequenced in four runs by a NovaSeq 6000 sequencer (Illumina) to generate paired-end 100bp reads. The raw FASTQ files from four flow cells were combined for each sample before downstream processing. RNA-seq data were analyzed using a local Galaxy instance (*52*). Quality and adapter trimming were applied to the raw sequencing reads using the Trim Galore! Package (Felix Krueger, github.com/FelixKrueger/TrimGalore). The reads were mapped to the human genome (UCSC hg19 with GENCODE annotation) using RNA STAR (*39, 40*). Per gene read counts from each sample were generated by featureCounts (*41*). The raw read counts were normalized and analyzed for differential expression between control and KMT5C overexpression tumors using DESeq2 (*53*). Genes with significant differential expressions were determined using an FDR-corrected P-value cutoff of 0.1. Significant genes and their fold-changes were analyzed by Ingenuity Pathway Analysis (IPA) software (Qiagen) to predict changes in Diseases and Biological Functions (*54*).

### KMT5C gene expression correlations with motility/invasion and OXPHOS genes across multiple tumor types

Co-expression data for KMT5C from the following tumor types were downloaded from cBioPortal: kidney renal clear cell carcinoma, thyroid carcinoma, uterine corpus endometrial carcinoma, bladder urothelial carcinoma, breast invasive carcinoma, head and neck squamous cell carcinoma, prostate adenocarcinoma, skin cutaneous melanoma, pancreatic adenocarcinoma, thymoma, and testicular germ cell cancer (*48*). All co-expression studies were based on TCGA’s patient RNA-seq data (Firehose Legacy, mRNA expression RNA Seq V2 RSEM) (*33*). Motility/invasion and OXPHOS gene signatures related to metastases were obtained from previous studies, and expression correlations to KMT5C were analyzed (*13, 31*). Positive or negative correlations between KMT5C and metastasis-related genes were determined using Spearman’s correlation coefficient cutoff of +/- 0.2. Correlation coefficients between −0.2 and 0.2 were not shown (marked gray in the heatmap).

### Patient gene co-expression analyses of KMT5C, motility, and OXPHOS genes in breast cancer and pancreatic cancer

Breast cancer or pancreatic cancer mRNA expression data on TCGA were downloaded from cBioPortal (TCGA, Firehose Legacy, RNA Seq V2 RSEM) (*33, 48*). For each tumor type, mRNA expression Z-scores for KMT5C and other genes of interest were plotted for all patients as a heatmap using cBioPortal. Rows represent expression levels of KMT5C, motility/invasion genes, or OXPHOS genes from patient samples (*13, 31*). Columns represent individual patients grouped based on the relative expression level of KMT5C. KMT5C high or low expression was determined with a Z-score threshold of +/- 0.5.

### Pathway activities in TNBC patients with increased or decreased levels of BACH1, RKIP, or KMT5C

Breast cancer patients with high or low expression levels of KMT5C, RKIP, or BACH1 were stratified with a Z-score threshold of +/- 1, using TCGA patient data accessed through cBioPortal (TCGA, Firehose Legacy, RNA Seq V2 RSEM) (*33, 48*). TNBC patient sample IDs were identified using the patient clinical data downloaded from Genomic Data Commons Data Portal (*55*). Samples indicated the TNBC status with negative “ER status by IHC”, negative “PR status by IHC”, and at least one negative result from “HER2 status by IHC” or “HER2 FISH status”. RNA-seq RSEM normalized count data for TNBC samples were downloaded from UCSC Xena and used for differential expression analyses (*56, 57*). Differentially expressed genes in KMT5C-high over KMT5C-low patients, RKIP-high over RKIP-low patients, and BACH1-high over BACH1-low patients were determined using the Limma-Voom package via the Galaxy server (*52, 58, 59*). For Limma-Voom, patient samples’ RNA-seq data were supplied as a single count matrix, and high or low indicators for all samples were designated in a factor file. Differentially expressed genes were plotted as volcano plots for each comparison (*60*). IPA was used to predict activities of related canonical pathways based on relative expression levels of KMT5C, RKIP, or BACH1 (*54*). IPA-predicted canonical pathways activation Z-scores from each comparison (KMT5C-high over KMT5C-low, RKIP-high over RKIP-low, and BACH1-high over BACH1-low) were compared to each other to identify pathways with the biggest activity changes across every comparison. IPA used the following FDR-adjusted P-value (q-value) cutoffs for each dataset to perform the canonical pathways cross comparison: KMT5C high vs. low q < 0.01, RKIP high vs. low q < 0.05, BACH1 high vs. low q < 0.01. These cutoffs were implemented due to limitations set by the IPA software.

### KMT5C expression and patient survival

Breast cancer or pancreatic cancer patients were stratified based on relative expressions of KMT5C. For each tumor type, patients with high or low expression of KMT5C were selected using cBioPortal with a z-score threshold of +/- 1 from the TCGA Firehose Legacy RNA Seq V2 RSEM datasets (*33, 48*). Survival analyses were performed based on TCGA’s patient overall survival data, and Kaplan-Meier plots were generated for each tumor type using cBioPortal (*33, 48*). A statistically significant difference in patient survival was determined by the logrank test on KMT5C-high and KMT5C-low patients with a P-value below 0.05.

### Additional statistical analyses and software

GraphPad Prism version 9 (GraphPad Software) was used for unpaired t-tests as indicated in the figure legend, with significant P-values shown (significance is determined by P-value < 0.05). Error bars represent standard deviations. All FDR-corrected P-values and Z-scores utilized the Benjamini-Hochberg (BH) correction method (*61*). Figures were created with Prism, BioRender.com, or Morpheus (Broad Institute, software.broadinstitute.org/morpheus).

## Supplementary Note

Gene expression is often conceptualized as a Poisson process, for which the variance equals the mean:

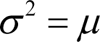

This implies that the CV^2^ and the Fano Factor *F* depend on the mRNA expression mean in the following ways:

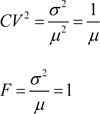

Thus, for a perfect Poisson process, the CV^2^ decreases as the inverse of the mean, while the Fano Factor is a constant = 1. However, many studies have indicated that eukaryotic gene expression can be non-Poisson, often due to intermittent bursts of gene expression (*7, 62*). Bursting modifies the above equations for mRNAs as follows.

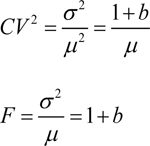

For mRNAs with zero-order production without bursts, at a constant rate *k* and first-order degradation at rate *g*, the mean and variance take the following values:

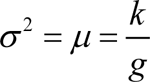

Therefore, the Fano Factor is =1 and the CV^2^ for such a process will be:

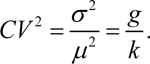

With bursting, these formulas become

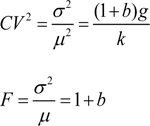

Typically, gene regulation affects the mRNA synthesis rate *k* rather than the degradation rate *g*. Assume that the synthesis rate of some mRNA increases by a factor *κ* due to RKIP overexpression.

The expected new mean, CV^2^ and Fano Factor will be:

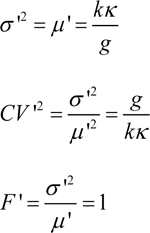

The corresponding ratios will be:

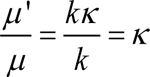

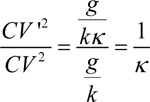

These ratios turn into differences of the corresponding log-ratios after taking the logarithm:

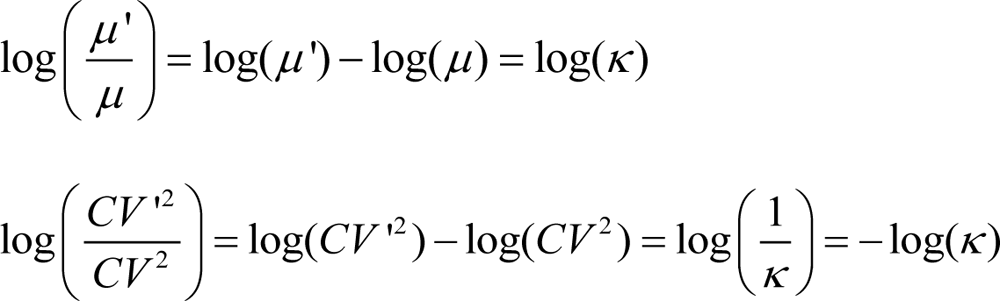

Therefore, plotting Δ log(*CV* ^2^) versus Δ log(*μ*) should result in a set of points that distribute along a line with slope = −1.

## Supplementary Materials

**Supplementary Figure 1.**
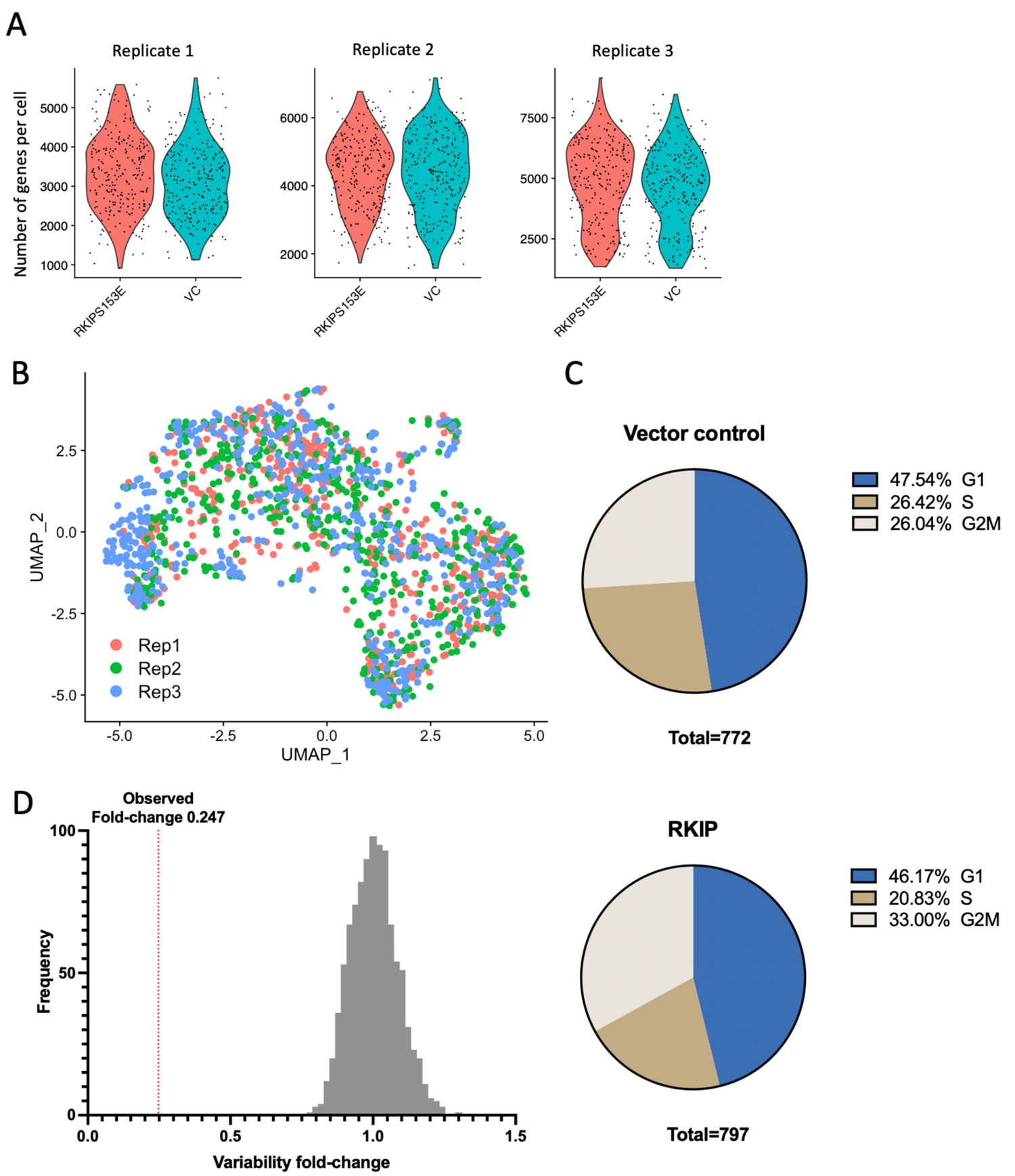
Results from BM1-CTRL and BM1-RKIP scRNA-seq. A) Number of genes mapped in each tumor cell in BM1-Ctrl or BM1-RKIP tumors from each replicate. B) UMAP clustering of single cells from each replicate. C) Distribution of cell cycles in BM1-Ctrl or BM1-RKIP cells. D) Observed fold-change of the overexpressed RKIP gene in BM1-RKIP cells compared to the distribution of results from 1000 permutations.

**Supplementary Figure 2.**
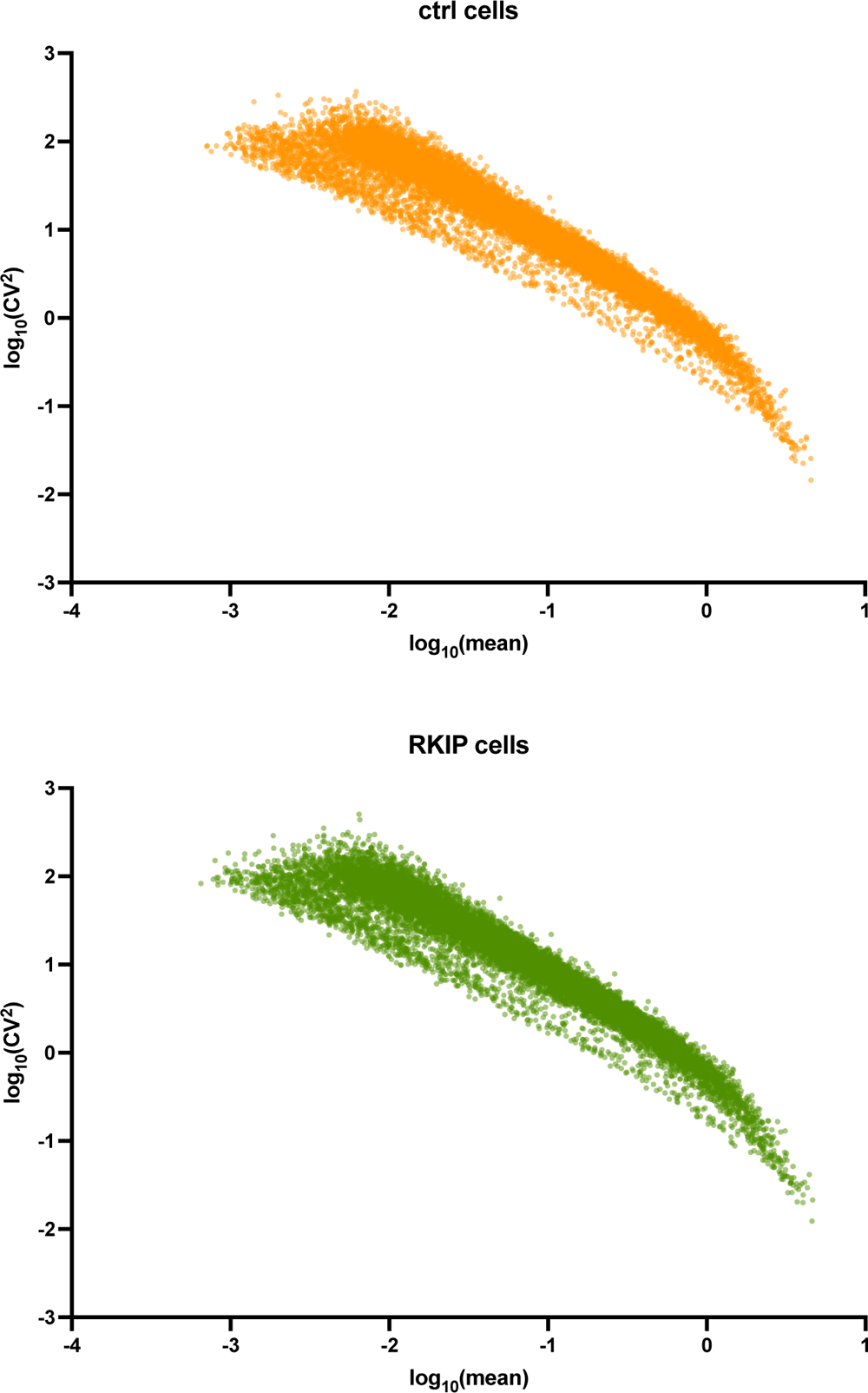
Gene expression mean and variability of BM1-Ctrl and BM1-RKIP cells. Gene expression log_10_(CV^2^) and log_10_(mean) were plotted for all genes in BM1-Ctrl cells (top) and BM1-RKIP cells (bottom).

**Supplementary Figure 3.**
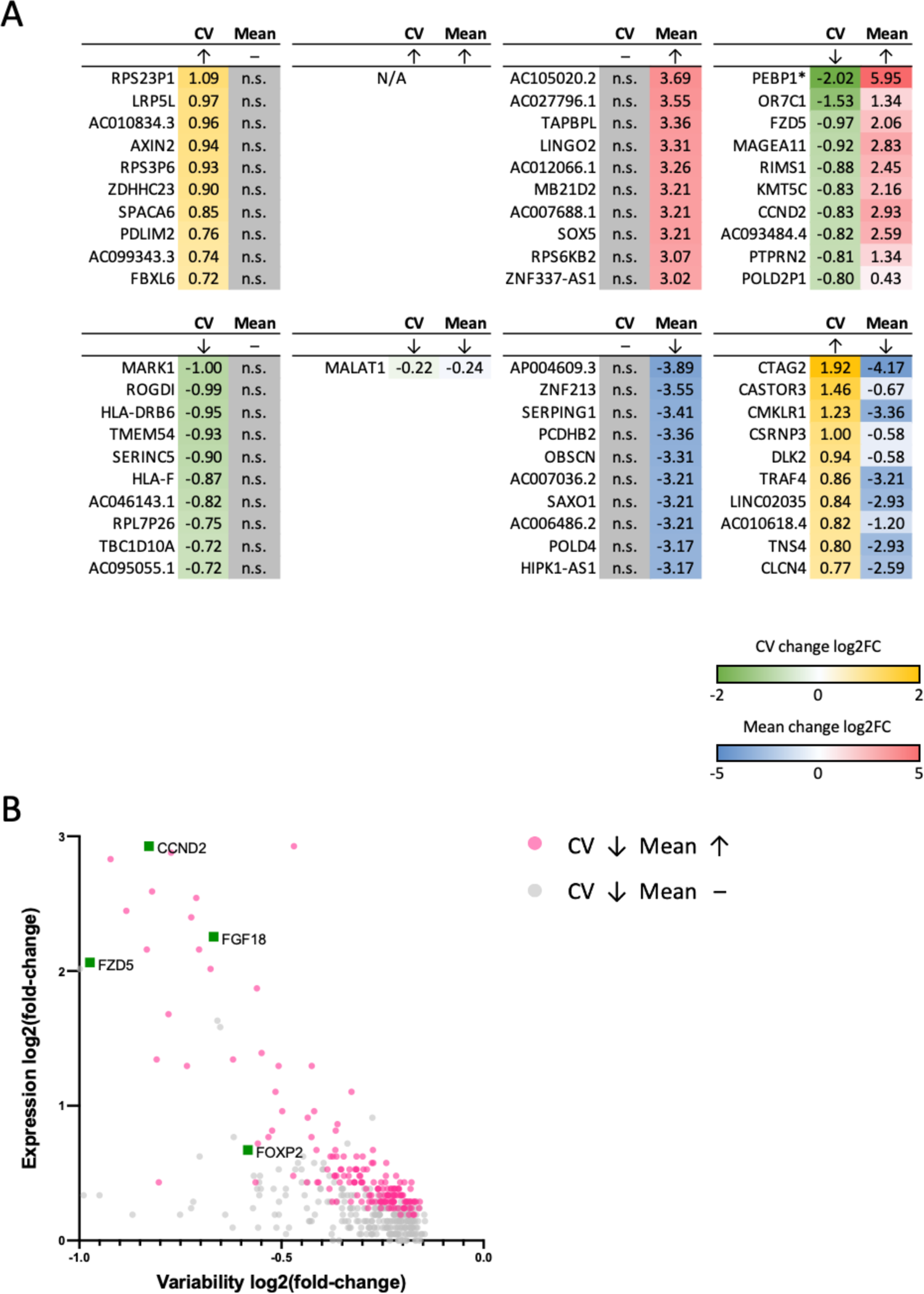
Genes with significant changes in mean or CV in BM1-RKIP cells versus BM1-Ctrl cells. A) Top genes from each direction of Fig. 2A. B) Genes with significantly less CV changes in BM1-RKIP cells are indicated with examples of cancer-related genes labeled.

**Supplementary Figure 4.**
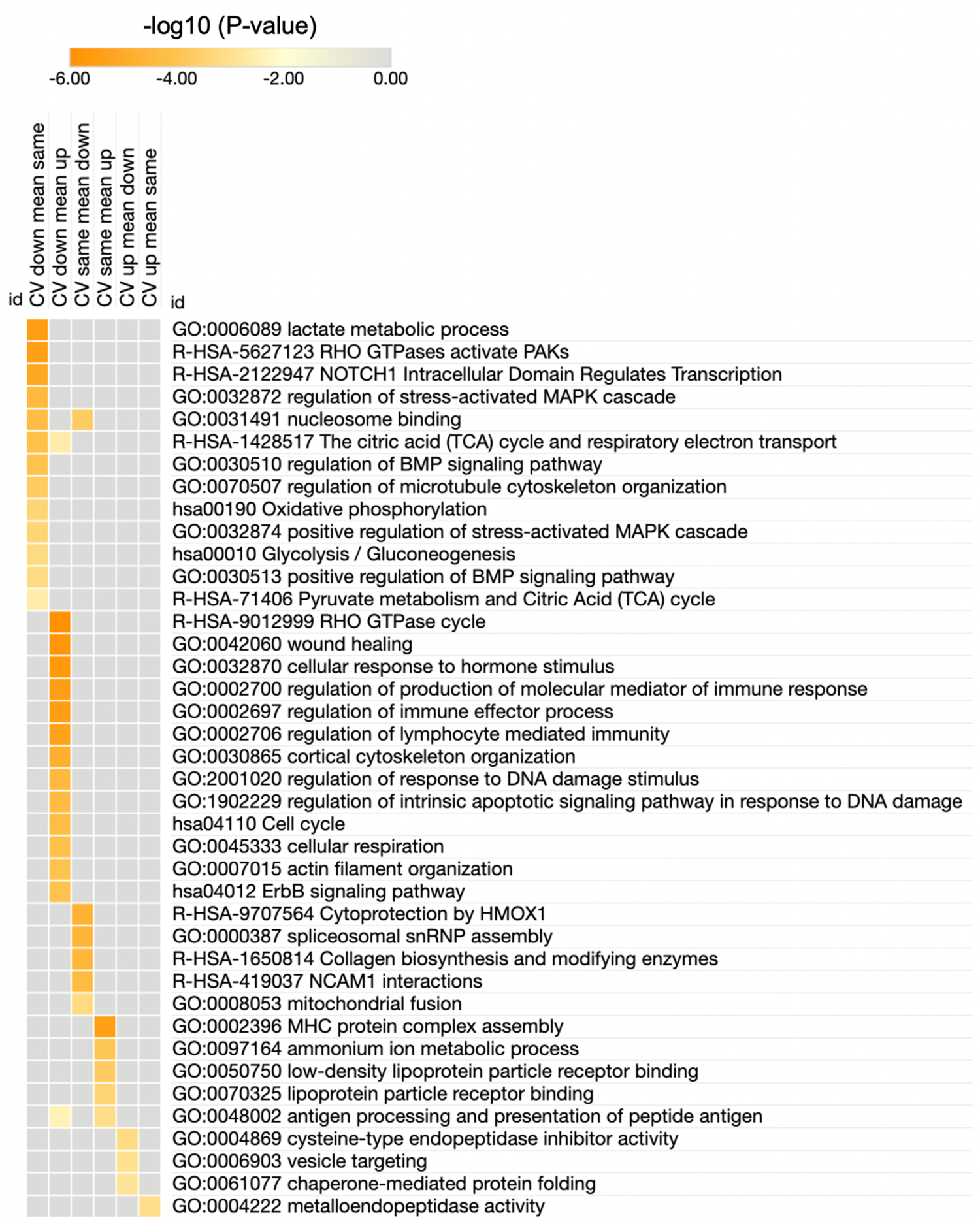
Significantly enriched gene sets from genes with changes in mean and/or CV in BM1-RKIP versus BM1-Ctrl cells.

**Supplementary Figure 5.**
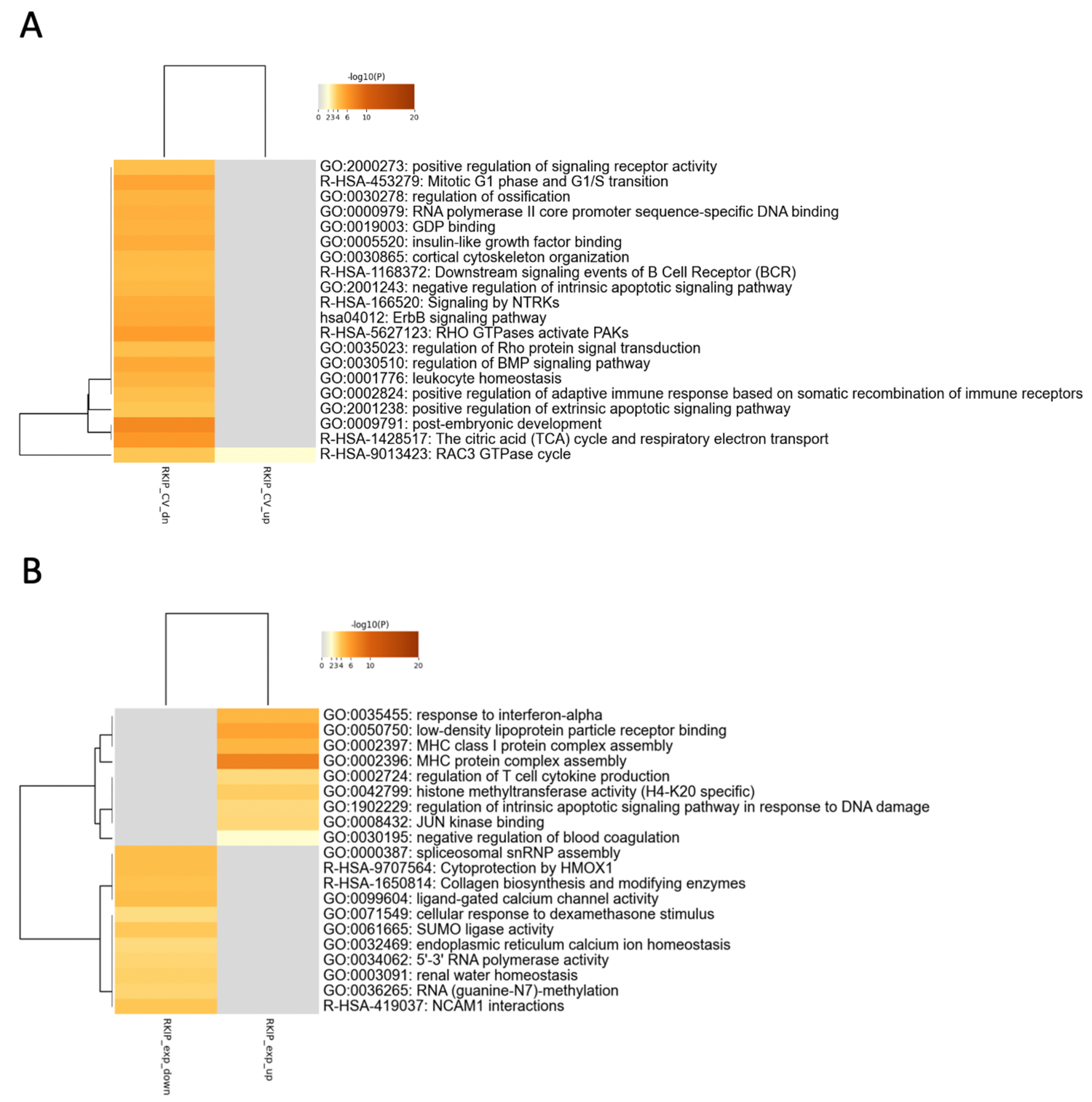
Significantly enriched gene sets from genes with changes in mean or CV in BM1-RKIP versus BM1-Ctrl cells. A) Enriched gene sets based on Metascape analysis of all genes with higher or lower CV in BM1-RKIP cells. B) Enriched gene sets from all genes up or down-regulated in BM1-RKIP cells.

**Supplementary Figure 6.**
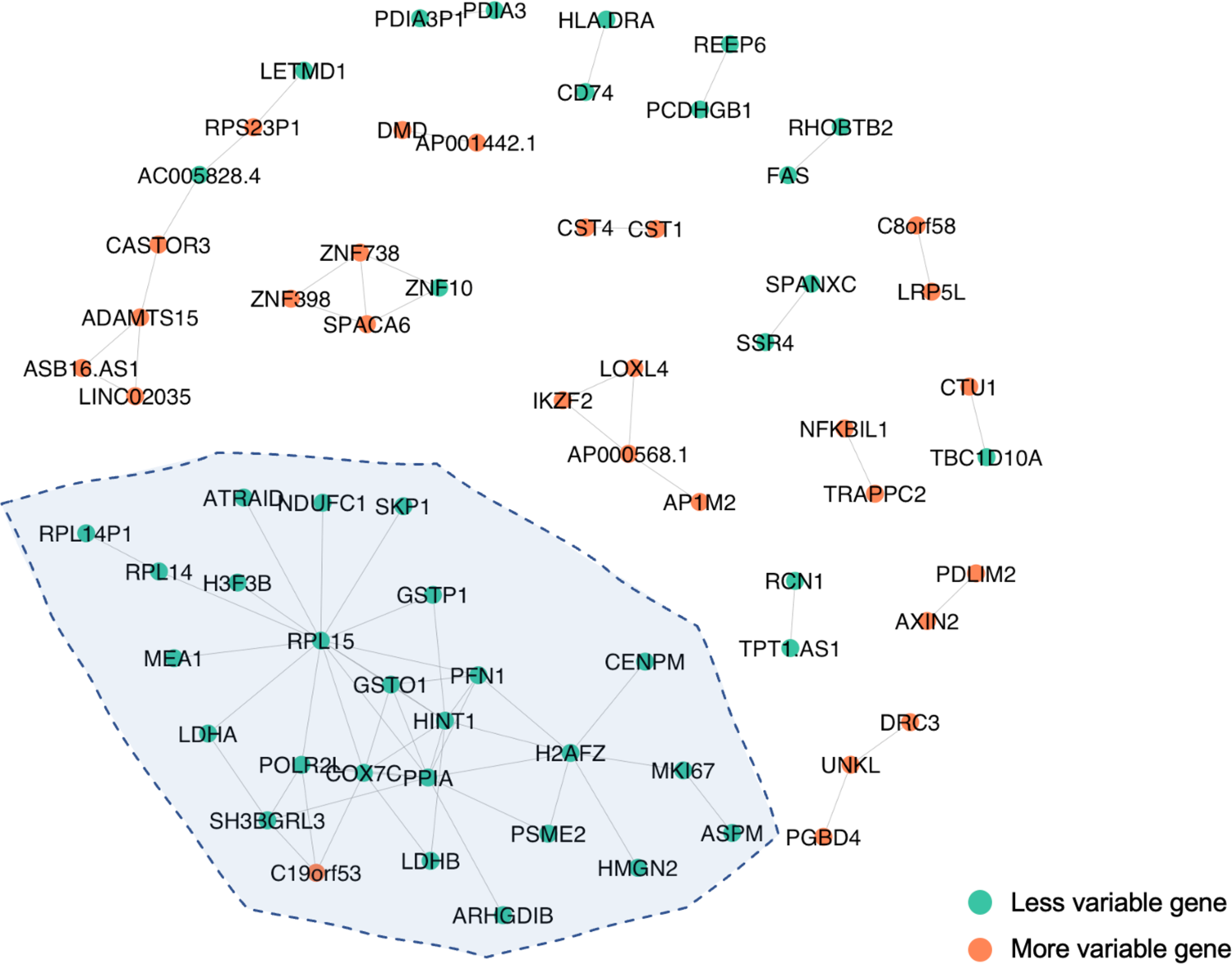
The complete gene co-expression network within BM1-RKIP cells from **Fig 3A**.

**Supplementary Figure 7.**
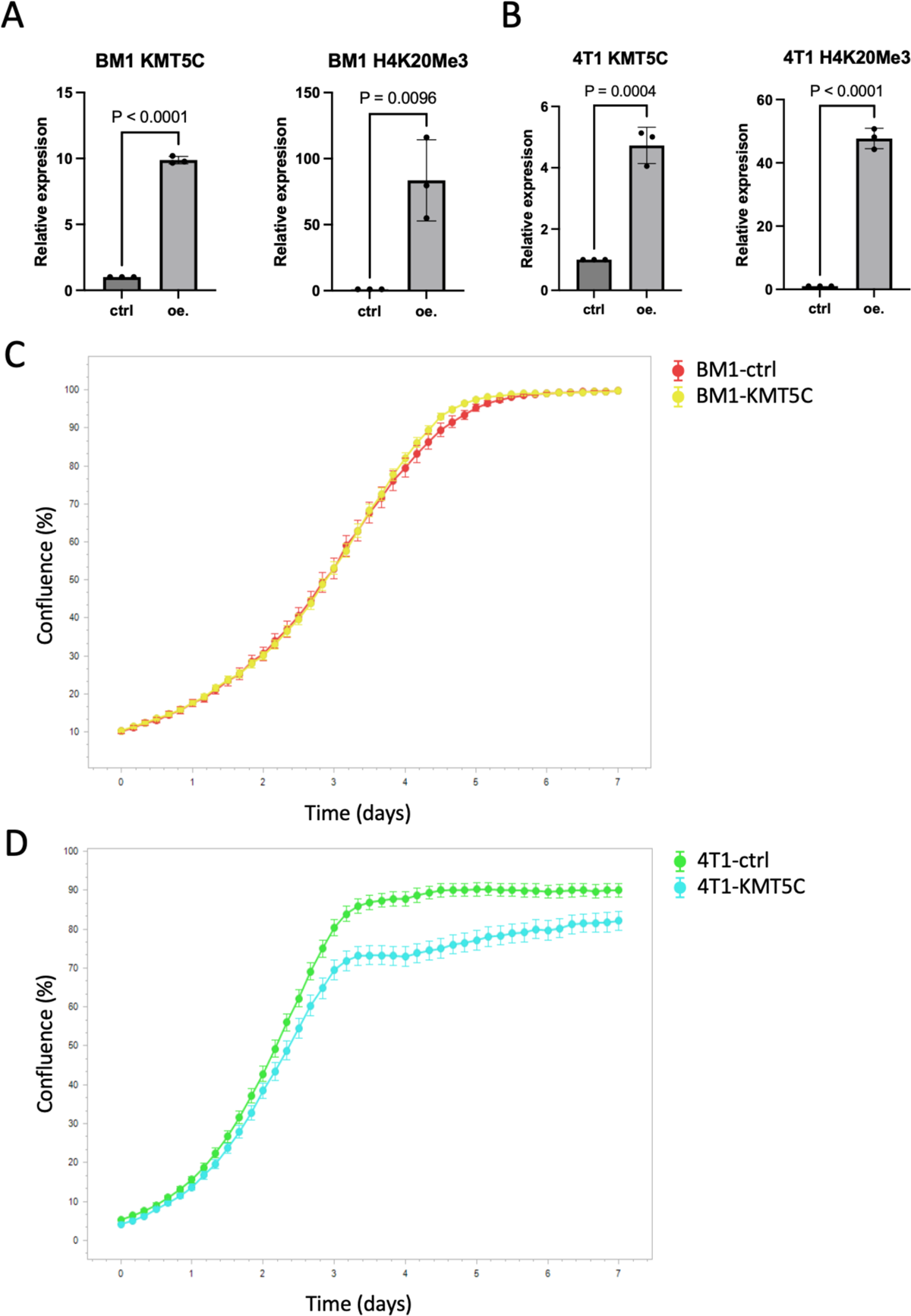
Validation of stable KMT5C expression in TNBC cell lines. A) Expression levels of KMT5C or H4K20Me3 in BM1 cells with (oe.) or without (ctrl) KMT5C expression. B) Expression levels of KMT5C or H4K20Me3 in 4T1 cells with (oe.) or without (ctrl) KMT5C expression. C) Proliferation curves of BM1-ctrl and BM1-KMT5C cells. D) Proliferation curves of 4T1-ctrl and 4T1-KMT5C cells.

**Supplementary Figure 8.**
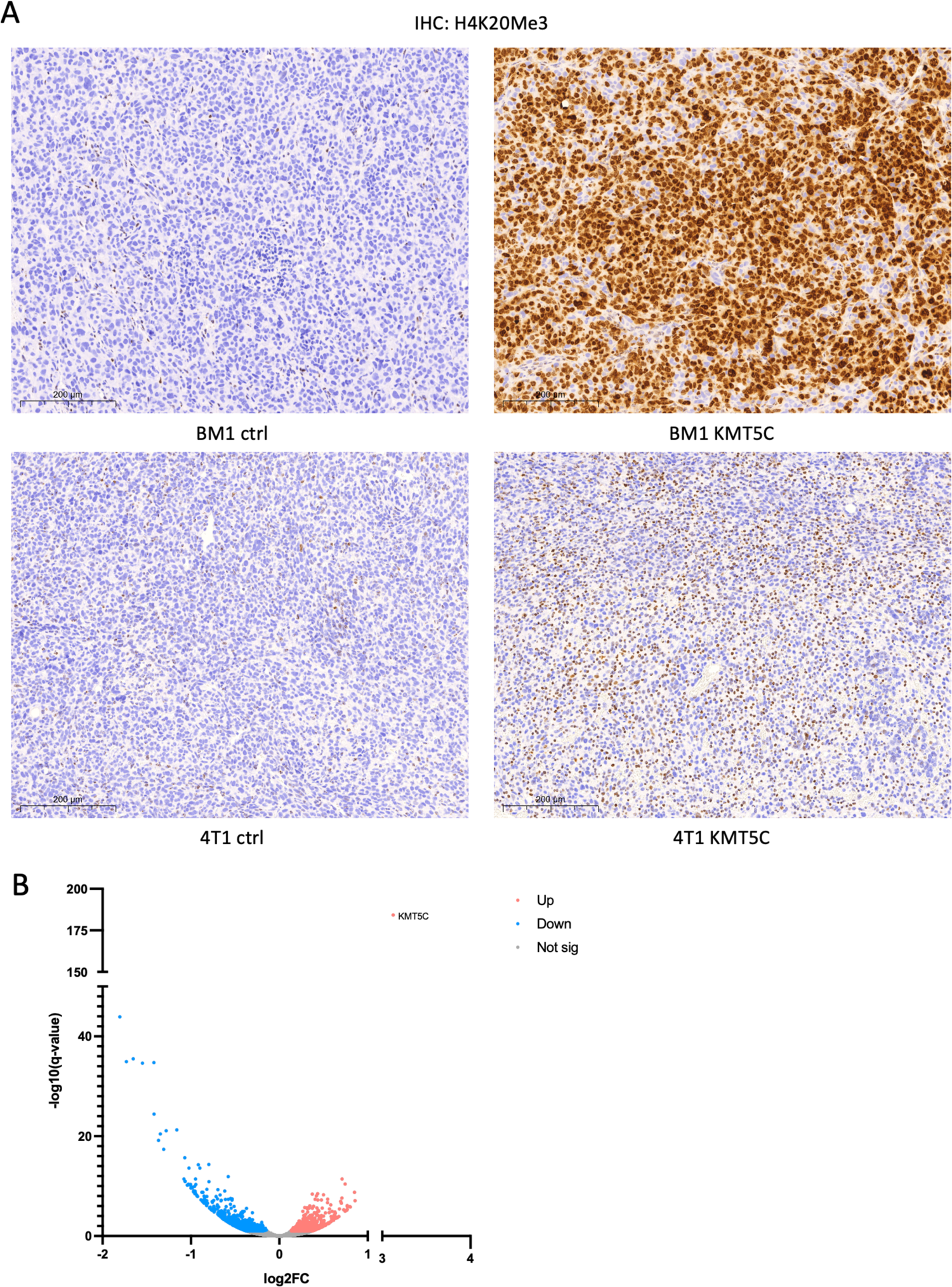
Effect of KMT5C overexpression in TNBC tumors in mice. A) IHC of H4K20Me3 in BM1 or 4T1 tumors with or without KMT5C expression. B) Volcano plot showing gene expression changes in BM1 tumors with KMT5C expression versus control tumors.

**Supplementary Figure 9.**
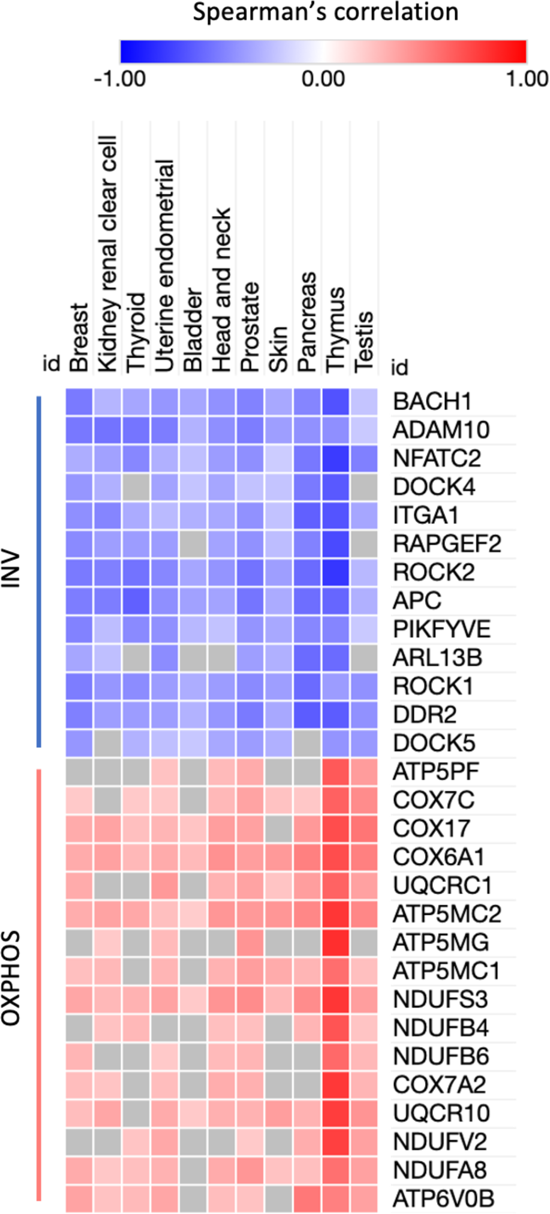
Spearman’s correlation between KMT5C and genes related to metastasis in cancer patients.

**Supplementary Figure 10.**
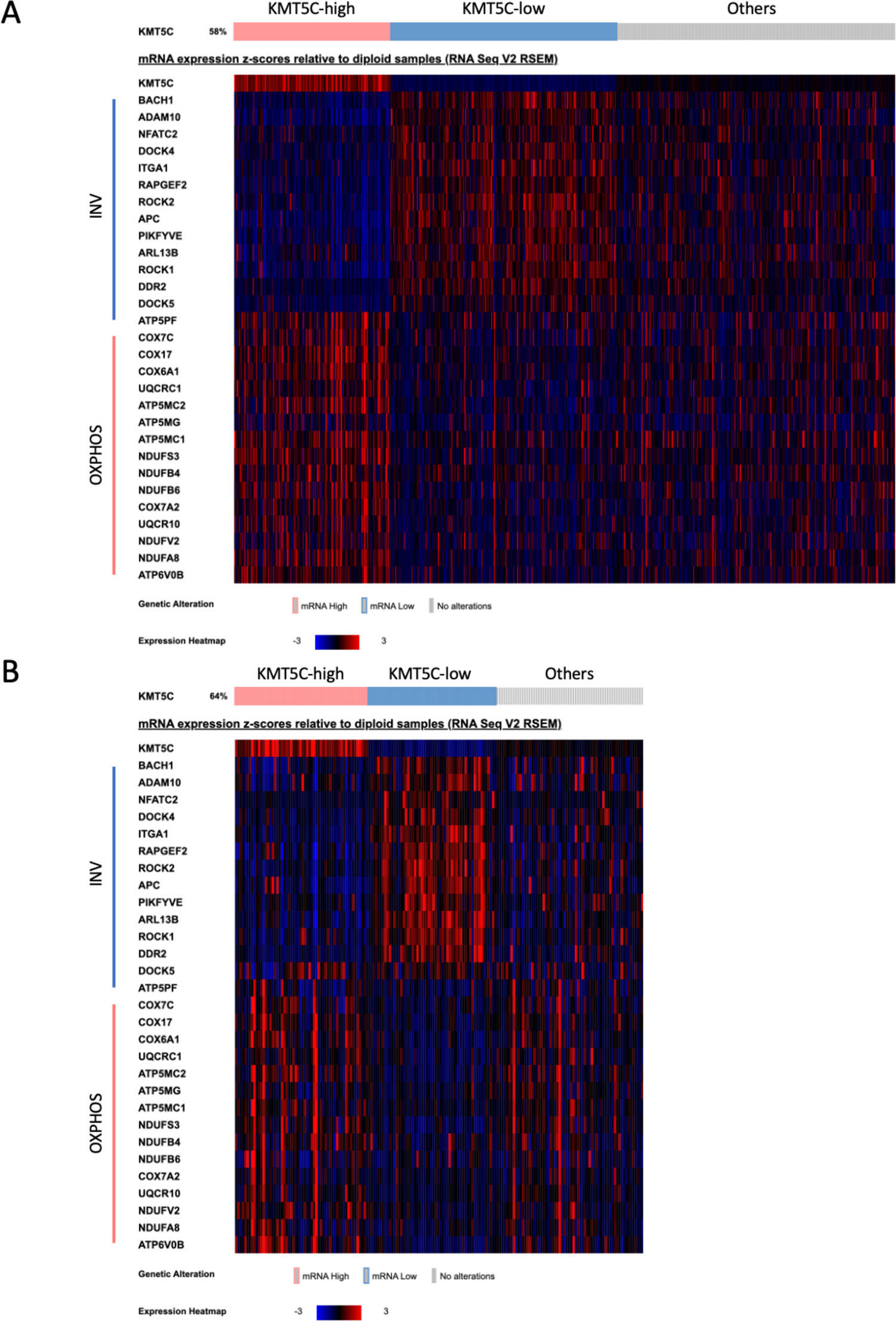
Relative expression levels of metastasis-related genes in breast cancer patients (A) or pancreatic patients (B) with higher or lower levels of KMT5C.

**Supplementary Figure 11.**
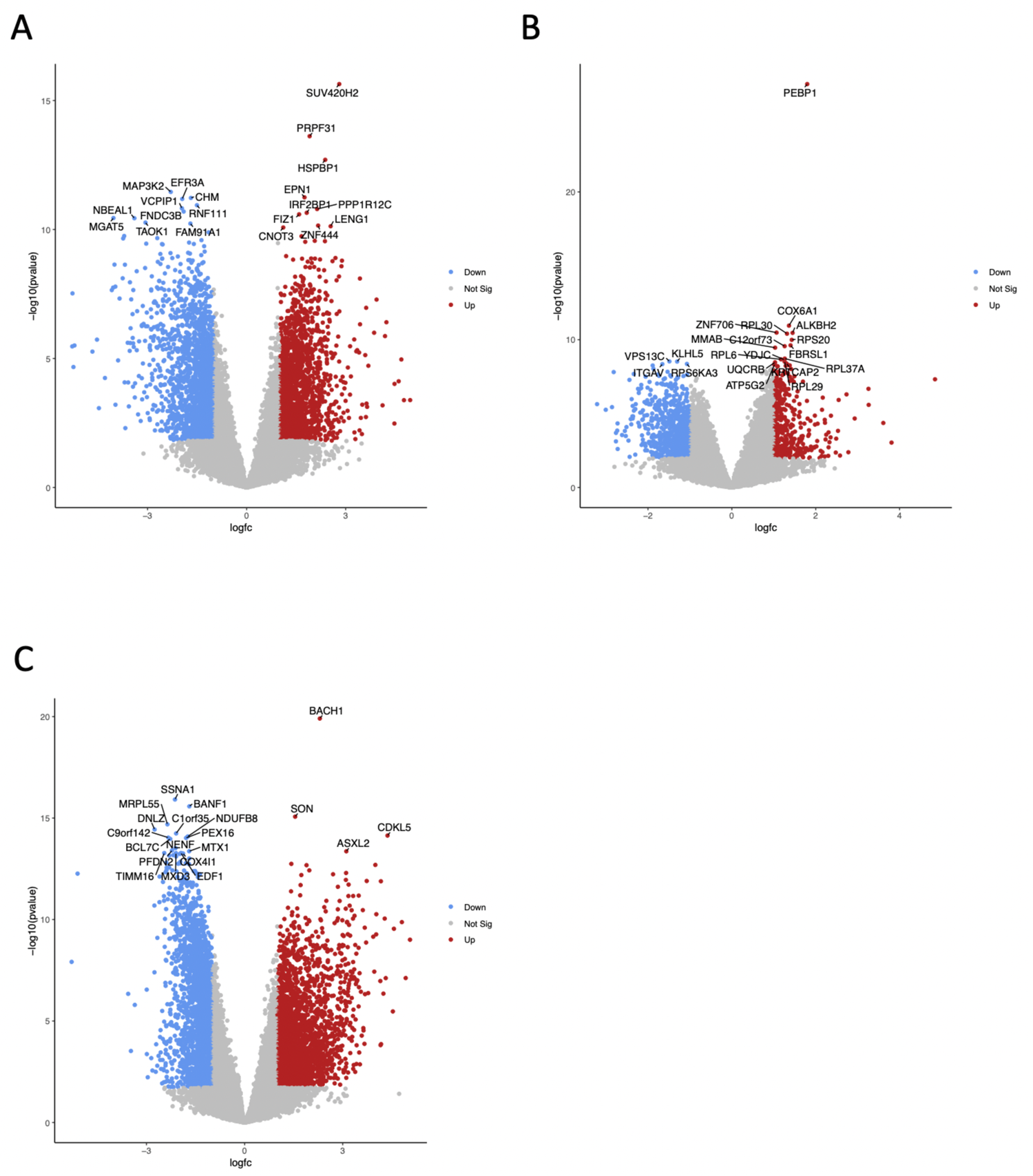
Gene differential expression analyses in TNBC patients. A) Volcano plot showing gene expression changes in TNBC tumors with higher versus lower levels of KMT5C. B) Volcano plot showing gene expression changes in TNBC tumors with higher versus lower levels of RKIP. C) Volcano plot showing gene expression changes in TNBC tumors with higher versus lower levels of BACH1.

**Supplementary Figure 12.**
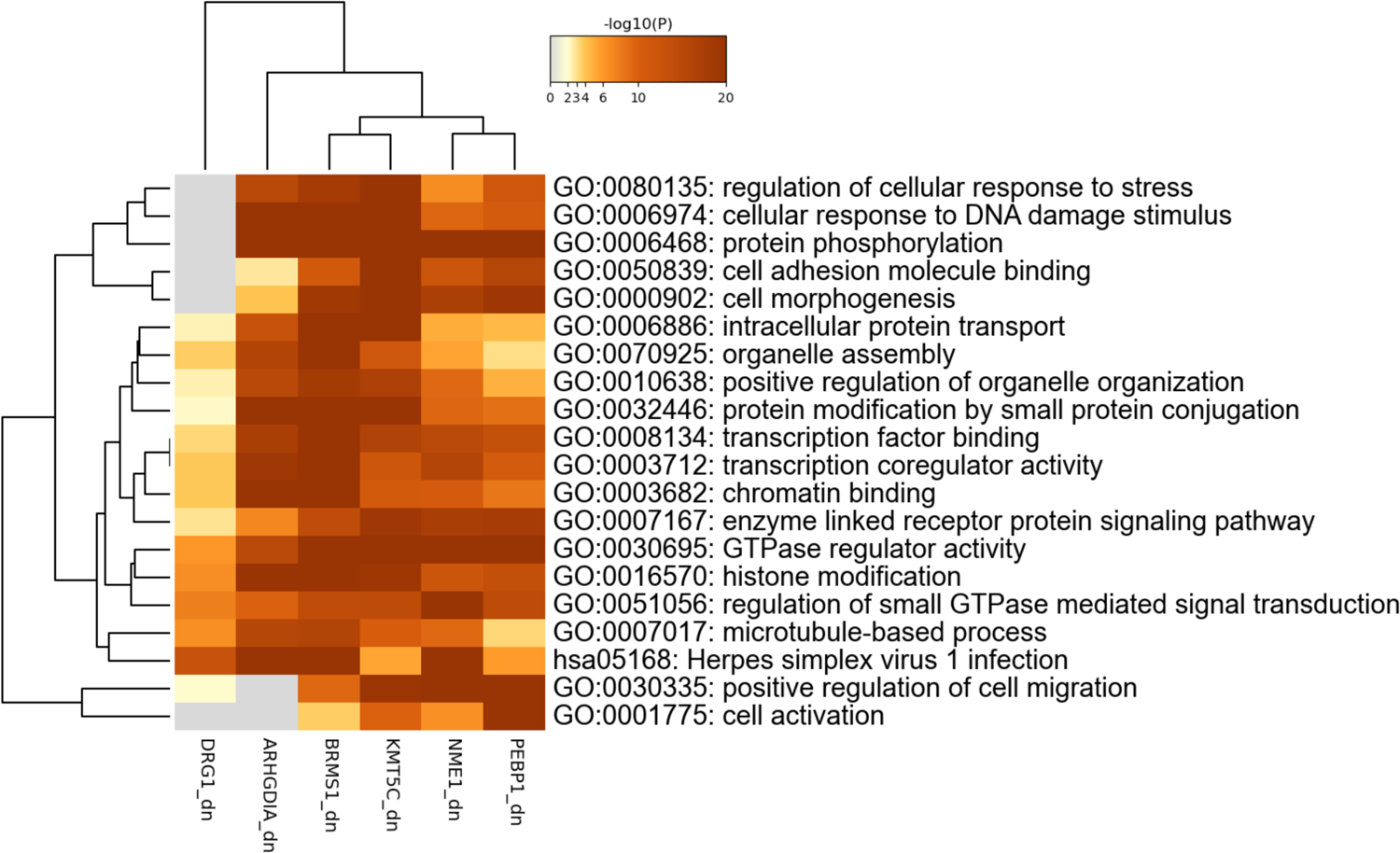
Significantly enriched gene sets from genes positively correlated with PEBP1, KMT5C, NME1, ARHGDIA, BRMS1, or DRG1 in breast cancer patients. Analyses of data from TCGA were done using Metascape.

**Supplementary Figure 13.**
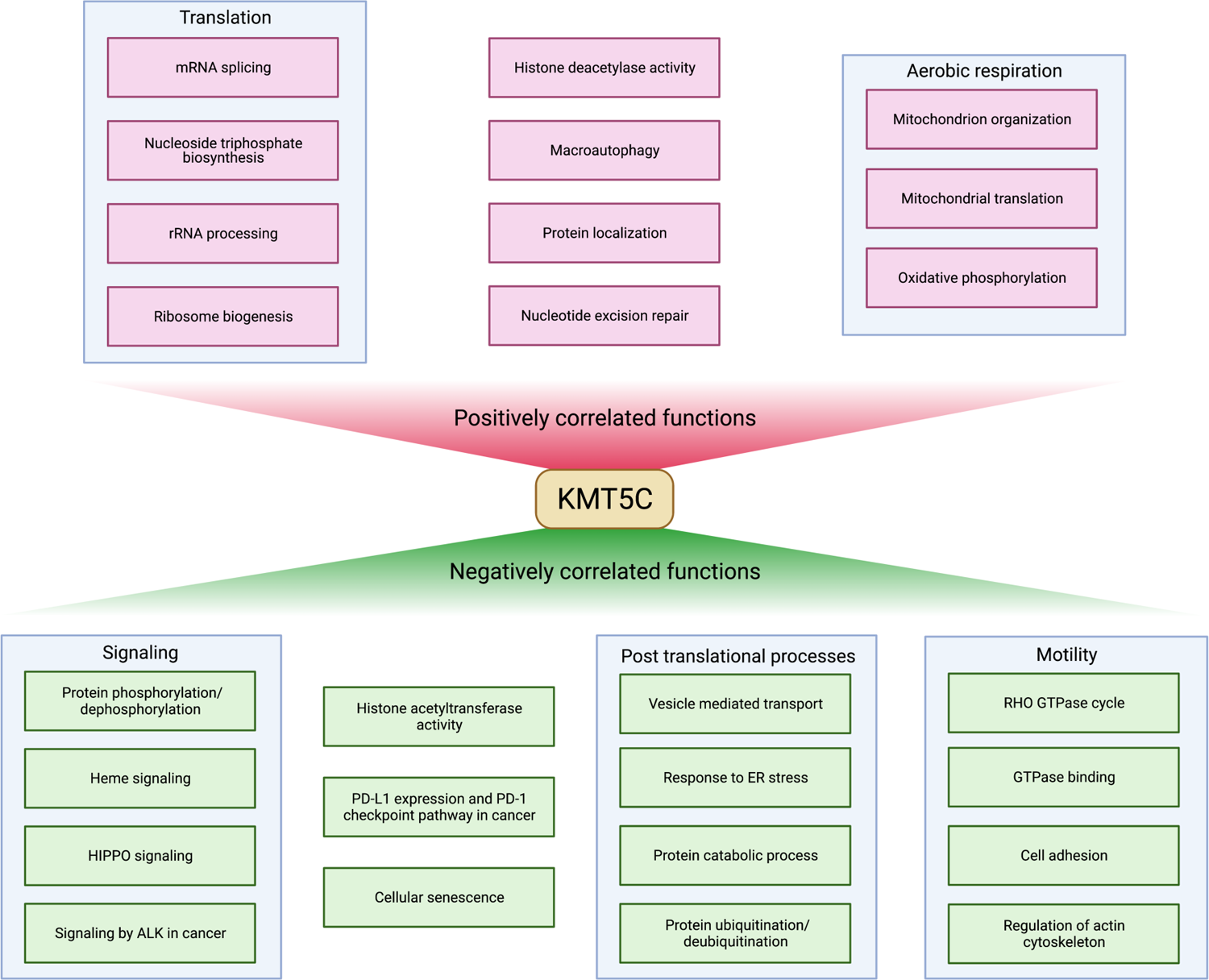
Significantly enriched gene sets positively or negatively correlated with KMT5C expression level in breast cancer patients.

## Data Availability statement

All data needed to evaluate the conclusions in the paper are present in the paper and/or the Supplementary Materials. Raw and processed RNA-seq data were deposited into the GEO database (GSE220671).

## Acknowledgements

We thank Sebastian Pott for valuable advice on experimental and computational methods, and Robert Rosner for helpful discussion of variability. We also thank Jonathan Willner for interpreting pathological features from IHC, Damian Berardi for advice on mouse models, and former and current members of the Rosner Lab (Ali Yesilkanal, Marcelo Fernandez De La Mora, Leticia Stock, Long Nguyen, and Margarite Matossian) for helpful feedback on the manuscript. We thank the University of Chicago Research Computing Center, Integrated Light Microscopy Core (Shirley Bond), Human Tissue Resource Center (Terri Li and Can Gong), Animal Resources Center (Ani Solanki), Genomics Facility (Pieter Faber), Cellular Screening Center, Center for Research Informatics, DNA Sequencing Facility, and Integrated Small Animal Imaging Research Resource for providing technical assistance. The results shown here are in whole or part based upon data generated by the TCGA Research Network: https://www.cancer.gov/tcga.

## Author Contributions

Conceptualization: MRR, DY Methodology: MRR, DY, GB, MC Software: DY

Formal Analysis: DY, GB Investigation: DY, CD, AV, LR-M, MH Resources: MRR

Data Curation: DY Writing: MRR, DY, GB Visualization: DY, GB, MC Supervision: MRR

Project Administration: MRR

Funding Acquisition: MRR

